# Utilizing a resource of enrichment profiles in plasma for the systematic assessment of antibody selectivity

**DOI:** 10.1101/158022

**Authors:** Claudia Fredolini, Sanna Byström, Laura Sanchez-Rivera, Marina Ioannou, Davide Tamburro, Rui M. Branca, Peter Nilsson, Janne Lehtiö, Jochen M. Schwenk

## Abstract

There is a strong need for procedures that enable context and application dependent validation of antibodies. Here we describe the foundation for a resource aiding more detailed assessment antibody selectivity for capturing endogenous proteins from human plasma. In 414 immunoprecipitation (IP) experiments with EDTA plasma, data was generated by mass spectrometry (LC-MS) with 157 antibodies (targeting 120 unique proteins). Out of a total of 1,313 unique proteins, 426 proteins (33%) were detected in > 20% of the assays and indicate a background comprised of mainly proteins from the complement system. For all proteins identified either in heat-treated or untreated EDTA plasma, frequencies of occurrence were derived. We determined z-scores for each IP as a measure of enrichment to annotate the antibodies into four categories (ON-target, CO-target, OFF-target and NO-target). For 45% (70/157) of the tested antibodies, the expected target proteins were enriched (z-score ≥3) above background. There were 84% (59/70) of binders that co-enriched other proteins beside the intended target, either due to OFF-target binding or predicted interactions. Comparing several antibodies raised against IGFBP2, the established library allowed us to describe protein complexes in plasma, and we employed multiplexed sandwich immunoassays to confirm these. In summary, the generated resource of plasma enrichment profiles and background proteins adds a very useful and yet lacking starting point for the assessment of antibody selectivity in this clinically important body fluid. The provided insights will contribute to a more informed use of validated affinity reagents and may lead to further advancements of plasma proteomics assays.

## MAIN TEXT

Antibodies are important tools used in a wide range of assays within life science, but there is a growing awareness about the importance to assess the quality of the data generated therewith [1]. To address this challenge, the recently formed International Working Group for Antibody Validation (IWGAV) proposed five strategies to assess the experimental performance of antibodies [2]. However, the opportunities for evaluating antibodies in the given context (= sample type) and application (= assays) are limited. In particular for body fluids there are currently no tools for modulating the samples to overexpress or delete expression of a target of interest. While the use of GWAS provides powerful opportunities to assess the relation between plasma protein abundance and genetic information [3, 4], affinity reagents are preferably still validated before the intended use. There is further need to experimentally assess selectivity of the affinity reagents and to enable the development of sensitive assays and technologies for proteins in plasma or serum.

Here, we aim to describe the foundation of a resource for the assessment of antibodies used to capture proteins from plasma. We utilize immunoprecipitation (IP) of full-length proteins in combination with mass spectrometry (MS) to systematically describe a library of identified proteins. In more than 400 IPs with plasma frequencies of occurrence (*f*) and enrichments scores (*z-scores*) were collected, and we list those proteins that were commonly detected as plasma background. Similar efforts have been applied to evaluate the performances of antibodies for IP in cell lysates [5]. Apart from few studies focused on specific, smaller number of targets [6, 7] however, there are no large scale and systematic studies extensively applying IP of endogenous full-length proteins and MS for antibody validation in plasma. Otherwise, peptide-specific antibodies to determine protein abundance after digestion of plasma proteins have been more frequently applied in combination with MS readout [8-10]. Additional approaches such as iMALDI [11] and MS-based immunoassays [12] complement activities using protein-enrichment before MS analysis. The composition of the plasma proteome was recently updated and now lists around 3,500 proteins detectable using MS techniques [13]. Adding further targets to this list, foster hence to perform deeper quantitative analysis of plasma proteins, and will require large recourses of validated antibodies. For MS-based techniques, indeed one of the keys is to use enrichment as a strategy to enhance the sensitivity [14], while purely affinity-based techniques will greatly benefit from the utility of highly selective binder in multiplexed assay systems [15].

### Study Overview

To establish a plasma-centric resource for selectivity analysis of antibodies, we applied a systematic approach using 157 antibodies (targeting 120 proteins) and a workflow depicted in **Fig. 1A**. The assays were built on a previously described procedure, in which antibodies are covalently coupled onto magnetic polystyrene beads prior to incubation with the sample [16]. The study included binders both of monoclonal and polyclonal origin and different species. In order to compare the performance of different antibodies raised against a common antigen, a subset of 25 proteins (21%) were targeted by more than one antibody (**Supplementary Fig. 1**). The selection of presented targets was driven by giving priority to antibodies raised against proteins known to be part of the plasma proteome and to be associated to a disease (**Supplementary Excel Table, sheet: “Selected Targets”**). As described in **Fig. 1B**, the majority of binders (65%, N=101) were raised against target proteins detected previously ‘in plasma’ (48% N=75) or annotated as ‘extracellular’ (17%, N=26), and fewer proteins were annotated as ‘cellular’ (36%, N=56). As reference for protein abundance in plasma, we considered the estimated concentrations found in the 2017 built of the plasma PeptideAtlas [13].

**Figure 1A:**
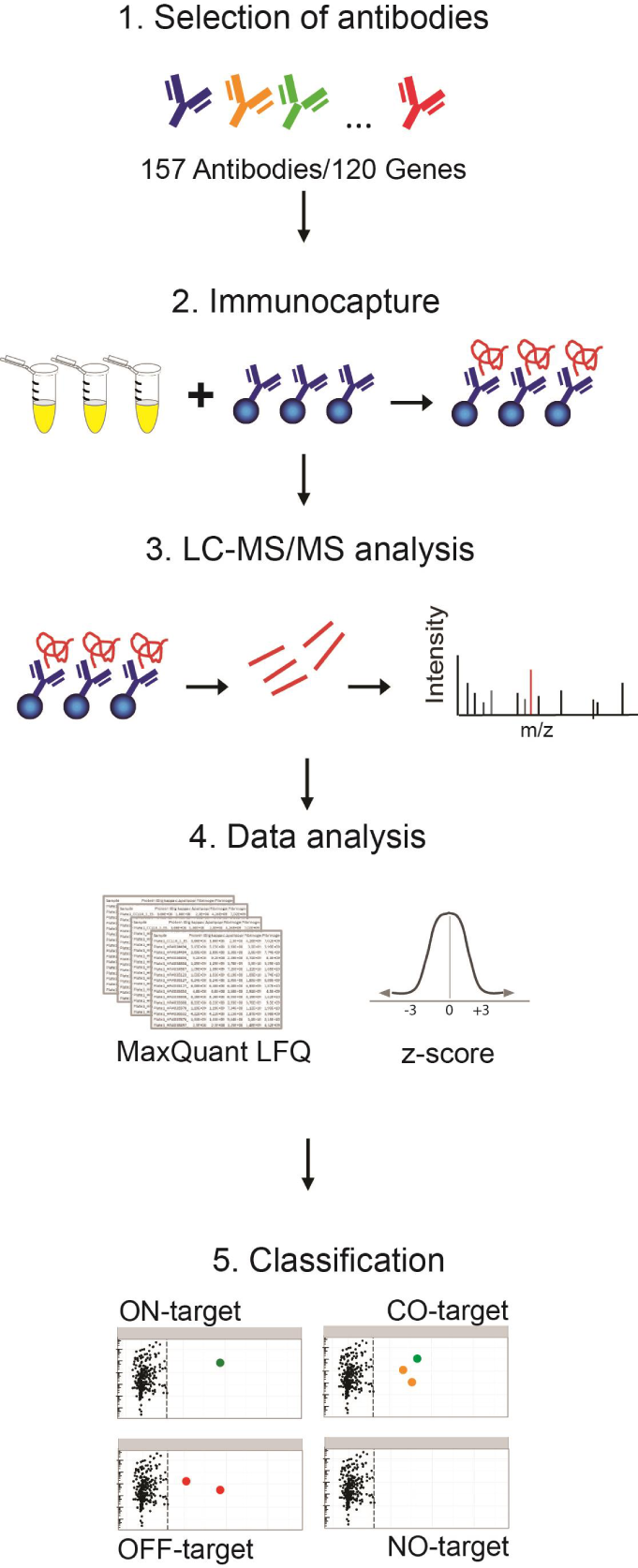
Workflow and study overview. A set of 157 antibodies targeting 120 genes were covalently coupled to magnetic beads and incubated one-by-one with EDTA plasma. Two to four replicate incubations were performed for each antibody. Following target enrichment, washing and digestion on beads, the obtained data files from LC-MS were searched and normalized by MaxLFQ, z-score analysis was performed to rank proteins specifically enriched by each antibody. Using the resource generated by > 400 IP assays, antibodies were classified based on their enrichment profiles: (1) ON-TARGET, only the target protein was enriched showing a z-score ≥3; (2) CO-TARGET, the target protein was enriched together with other proteins also associated to a z-score ≥3; (3) OFF-TARGET, only proteins other than the expected target were enriched; as well as (4) NO-TARGET, in case no protein was enriched (z-scores <3).

**Figure 1B:**
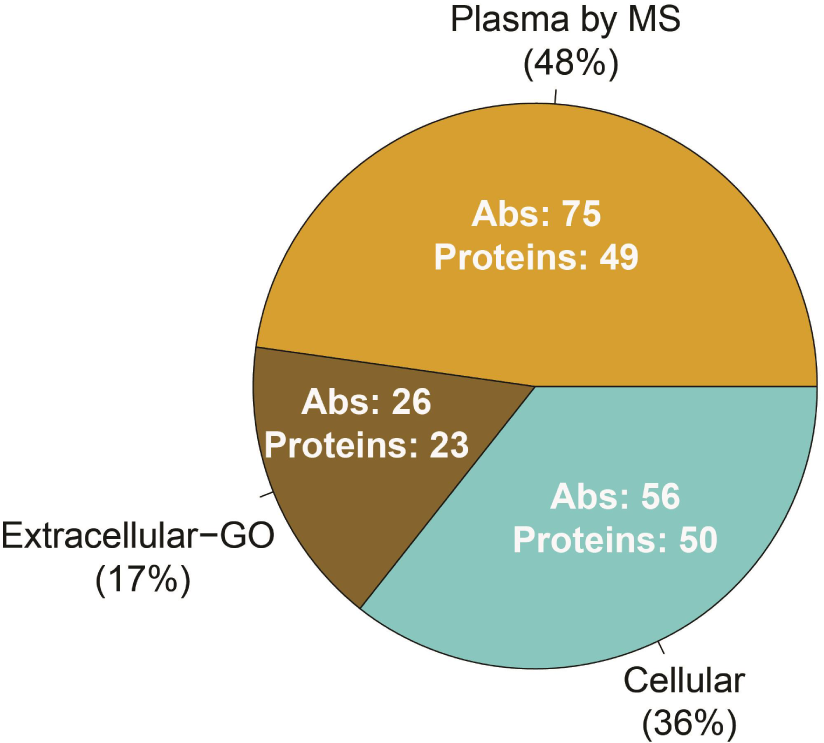
Distribution of antigen annotation. The target proteins of the 157 antibodies were grouped as: “Plasma by MS”, were identified in plasma previously by mass spectrometry as reported by Peptide Atlas. Cellular and Extracellular were assigned according to Gene Ontology classification (see Materials and Methods). Numbers stated inside the pie chart refer to the number of antibodies (Abs) in the category and corresponding number of target proteins.

**Figure 1C:**
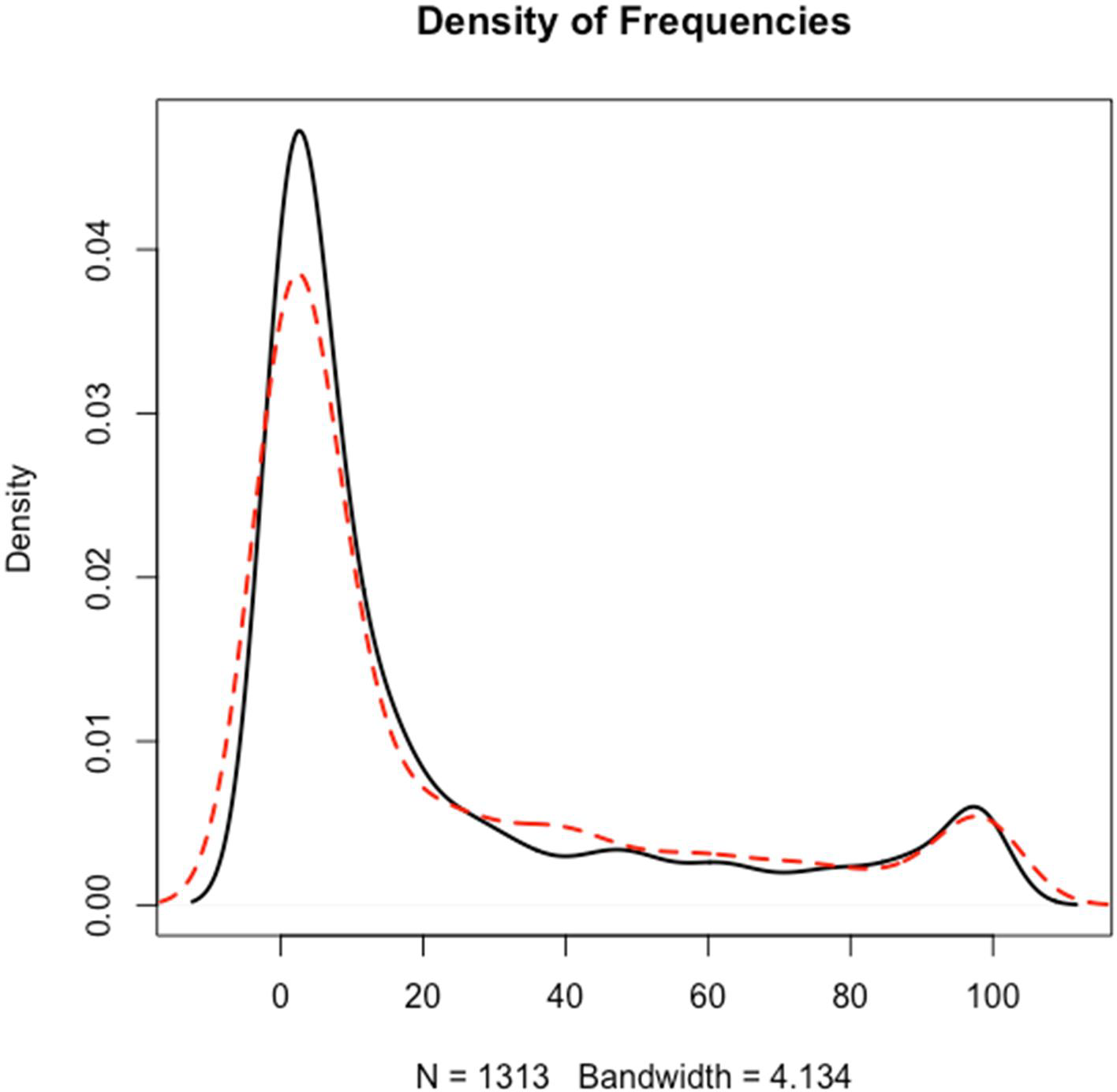
Distribution of frequencies of identification. The proteins obtained from assays conducted in heat-treated (red) vs untreated plasma (black) were collected in terms of the number of times they were observed in the IP-MS data. For both sample types, the majority of the 1313 proteins were found in less than 20% of the IPs.

**Figure 1D:**
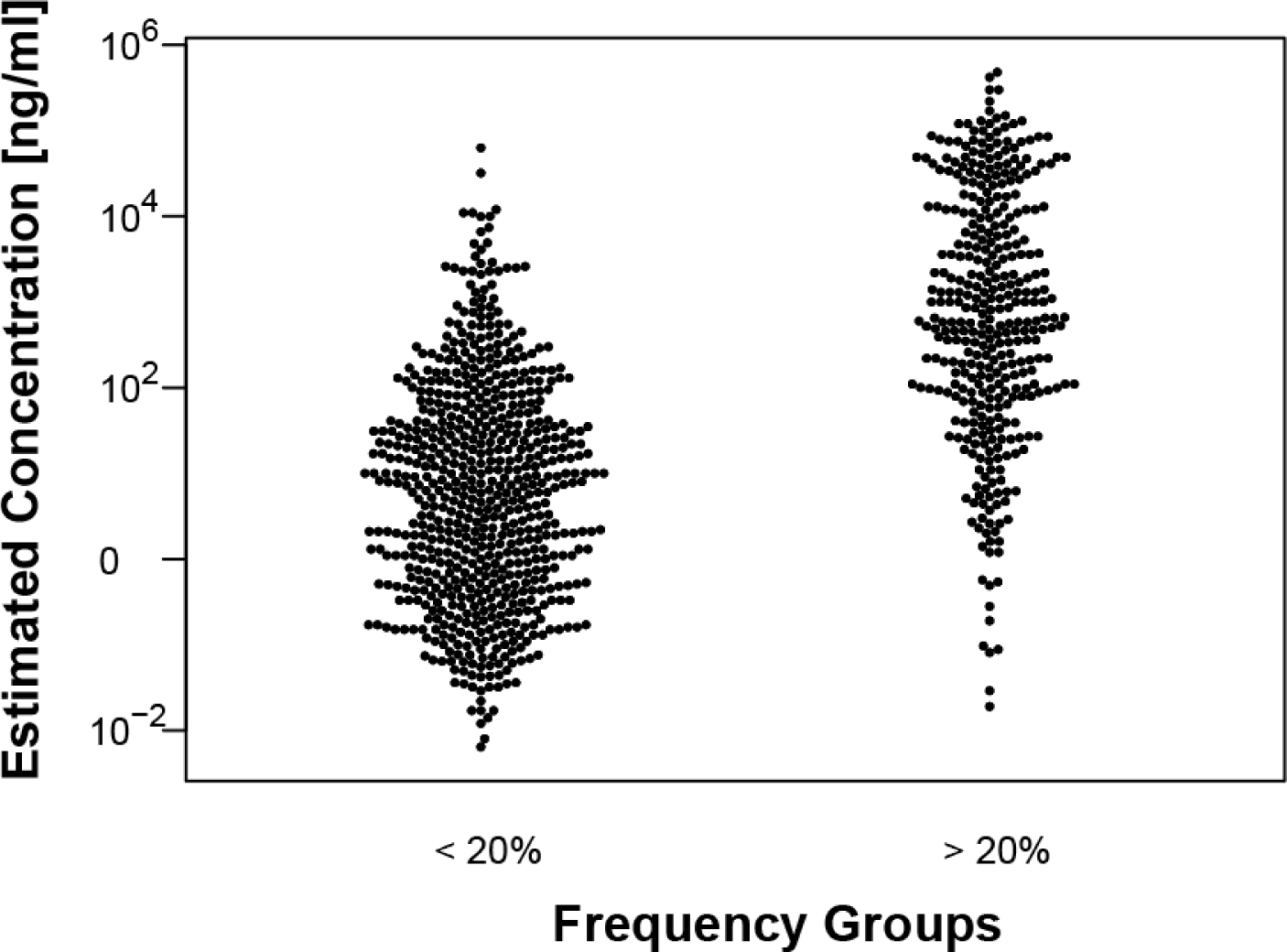
Frequency vs Concentration. Estimated concentrations in PeptideAtlas were compared between frequent (>20%) and less frequent (<20) protein identifications.

### Pilot study

In a pilot, we compared the experimental conditions to reach an analytical sensitivity to detect plasma proteins and to assess the potential to discriminate between specifically captured or frequently observed proteins (denoted plasma background). A set of antibodies obtained from commercially available ELISA kits for plasma protein analysis were used to cover a plasma concentration range between μg/ml (CRP) to low pg/ml (IL1A, see **Supplementary Note 1**). We tested combinations of different amounts of plasma (30-1000 μl) as well as increasing amounts of beads (30,000 - 1,000,000) to assess the detectability of the targets **Supplementary Fig. 2A**. We concluded that 100 μl of plasma and 500,000 beads were suitable for larger scaled investigations, enabling the detection within μg/ml and high pg/mL (e.g. KLK3 was detected in a pool of plasma obtained by mixing samples from healthy females and males, see **Supplementary Fig.2B**). For the detection of pg/ml plasma proteins, such as IL1A, the IP may require even larger amounts of sample and beads, which was found less suitable for large scale efforts (**Supplementary Fig. 2A**).The LFQ intensities from the IPs were used to calculate z-scores and compare peptides from replicated assays. In the pilot, all the antibody profiles barely reached an enrichment threshold (z-score ≥ 3) when using only this limited set of assays (*f*) in the library for data analysis. Utilizing a larger library of plasma IP data, as introduced and described below, improved the number of expected target proteins with z-scores ≥ 3 (**Supplementary Fig 2B**).

**Figure 2:**
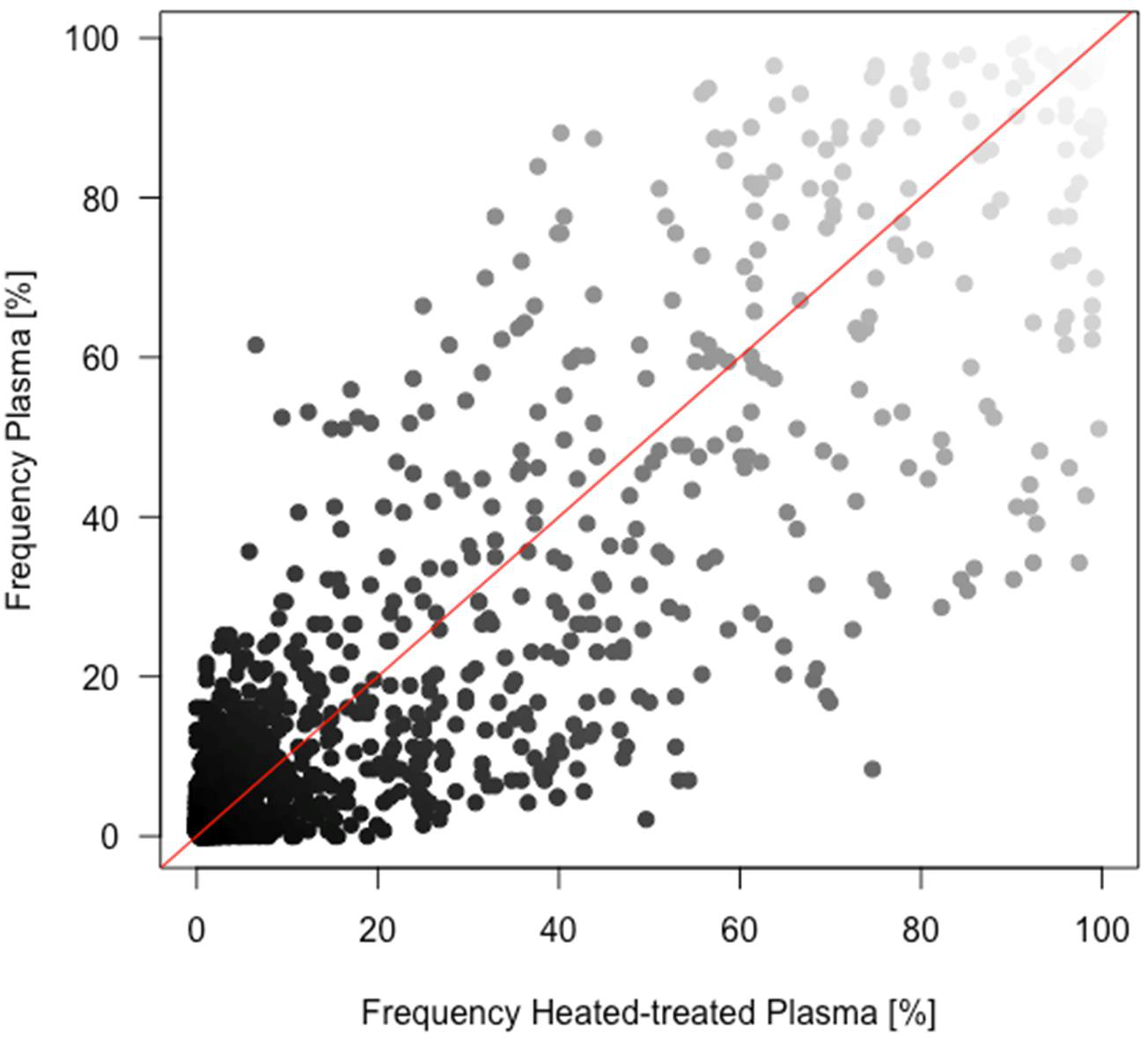
Frequency correlation. The relation between the frequency percentages of protein occurrence in IPs performed with heat-treated and untreated plasma is shown. The red line represents the line of identity.

### Assessing the resource

It is well accepted that MS provides in-depth information about the protein content of a sample, however hundreds if not thousands of proteins can be identified in a single IP experiment [17]. This calls for a careful assessment and interpretation of data from IP assays where many other proteins than the intended target can be identified in the same range of spectral counts or precursor intensities. As described in the context of cell lysates, the necessity to compare the outcome of several experiments, including negative controls or unrelated antibodies is essential [5, 18, 19]. Mellacheruvu and colleagues described how lists of background proteins that were obtained from control assays performed in similar experimental conditions can be used to assess specific enrichments in IPs [18]. In our case, all the data (not exclusively control IgG) was generated in separate experiments (refer to as “experimental batches” or “batches”). These sets of data were used to build a larger library of proteins and annotate these by their frequency of identification (*f*) and enrichment scores (z). These then serves as a resource to assess enrichment profiles of the antibodies. Details on experimental batches and frequencies for proteins are described in **Supplementary Excel Sheets “Experimental Batches” and “Frequencies of identification”**.

We applied MaxLFQ label free quantification to 414 plasma IPs (pIP), a total of 1,313 unique proteins were identified, excluding variable domains of immunoglobulin for heavy and light chains. The complete list of proteins can be found in **Supplementary Excel Sheet: “Frequencies of identification”.** Per IP, about 290 proteins were detected on average. Data was collected from experiments performed under heat treatment versus not heat conditions, as further discussed below, we found that the average number of identifications per experiment were slightly higher in heat-treated plasma (300±97 in 276 pIPs) compared to untreated plasma (283±97 in 138 pIPs). For both sample types, the majority of proteins (66%) were detected in *f* < 20% of all assays (**Fig 1C** and **Supplementary Excel Sheet: ”Frequencies of identification”**). Combining the frequencies of how many times a protein occurred in each IP, proteins found in *f* > 20% of all assays (denoted ‘frequent’) were compared with those found in *f* < 20% (denoted as ‘less frequent’). The resulting GO analysis revealed that terms related to the complement activation and wound healing (GO:0002576, GO:0006956, GO:0050817, GO:0009611, GO:0007596) were enriched for the frequent proteins (Supplementary Excel Sheet: ”GO enrichment analysis”). Other enriched terms for the frequent proteins were related to lipoprotein and their complexes (GO: 1990777, GO:0032994, GO:0034358) as well as vesicles (GO:0031983, GO:0060205). Further investigations also found a significant association (p-value <2*10^-16^) relation between the frequencies and estimated protein abundance in plasma [13], as well as to sequence coverage (**Fig 1D, Supplementary note 3** and **Supplementary Figure 3A-B**). Further details on frequency of identification and intensities variates along the experimental batches can be found in the **Supplementary Figure 4A-B** and **Supplementary Excel Sheet:’’Batches Kruskal Wallis test”**.

**Figure 3 A:**
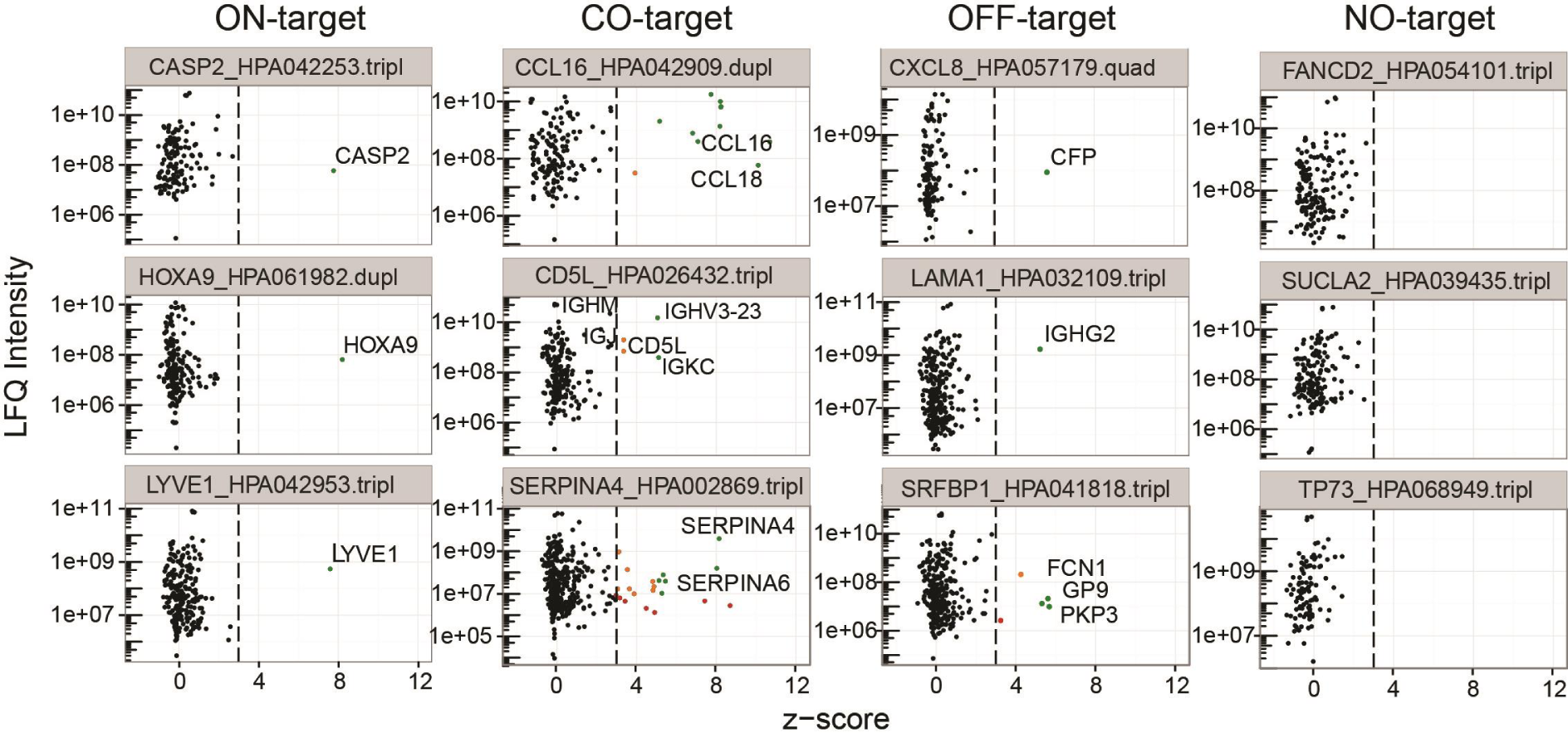
Classification of antibodies. Three representative examples of are given for each of the enrichment categories (ON-target, CO-target, OFF-target, and NO-target). On the top of each plot are the target gene, antibody ID and the number of replicates IP performed for the antibody. The dots in each plot represent identifications present in all the replicates available for the specific antibody.Green dots: z-score > 5 and LFQ intensity > 1e+07; yellow dots: z-score > 3 and LFQ intensity > 1e+07; red dots: z-score > 3 and LFQ intensity < 1e+07. Text: expected target and hypothesized off-targets or interactors. A complete list protein identified and relative z-scores are available in **Supplementary Excel Table**, Sheet **“z-score> 2.5”**.

### Differences between untreated and heat-treated plasma

Previously, we have shown that after heat treatment of plasma samples at 56°C, improved the limit of detection in plasma profiling assays using antibody bead array [20]. Heat treatment of plasma may indeed retrieve epitopes and making proteins more accessible to antibody binding, however heat may also affect the sample’s composition by causing aggregation and proteins to precipitate. As shown for all IPs in **Fig. 1C** and **Supplementary Figure 4A**, heat treatment may influence which proteins are listed among the common contaminants and their frequencies. When we consider LFQ intensities instead of frequencies, we observed that in heat-treated samples particularly fibrinogens (FGA, FGB and FGG) were more abundant (**Supplementary Fig. 5**). Previous observations state that denaturation of fibrinogen starts at 55°C (two transitions peaks at 57.7°C and 96°C) and particularly affects the D fragment [21]. In heat-treated samples, the affinity of fibrinogens to plastic surfaces has also been reported to increase [22]. Via a similar mechanism, fibrinogen’s unspecific binding to surfaces of magnetic beads or to heavy chains of IgG antibodies on the beads may be enhanced [23, 24]. Hence, heat-induced denaturation of fibrinogen may alter the frequency with which proteins of this family are identified. Other plasma proteins such as albumin, apolipoproteins (APOB, APOC2, APO2, APOE), Keratines (KRT1, KRT2, KRT10), Fibulin, Fibronectin 1 or IgM were less frequent among the common contaminants (see also **Supplementary Excel Sheet: “Frequencies of identification”**).

### Classification

Extending the use of the library containing common contaminants and frequencies, we aimed at classifying the antibodies based on their enrichment profiles in plasma. Considering a protein as being enriched when the LFQ intensity was z-score ≥ 3, all antibodies were annotated according to the following categories (**Fig. 3A**):

- ON-target: when z ≥ 3 was only assigned to the expected target.
- CO-target: other proteins besides the expected targets were detected with z ≥ 3. Here we will discuss three sub-categories related to the origin go the additional target.
- OFF-target: proteins other than the expected targets were enriched with z ≥ 3.
- NO-target: all detected proteins were classified with z < 3.

**Figure 3 B:**
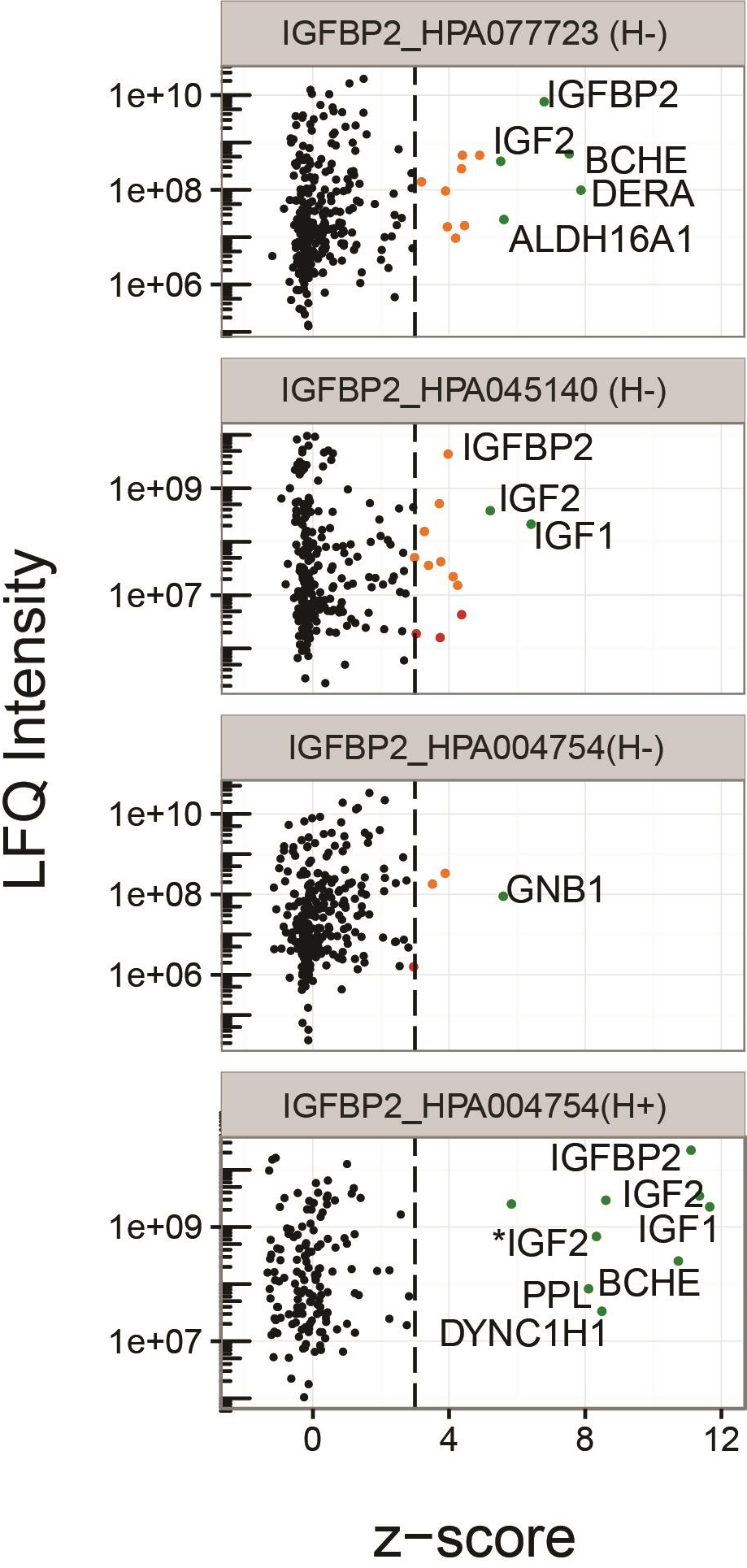
Paired antibodies and co-enrichment profiles. The z-score/LFQ intensity plots of paired antibodies raised against IGFBP2 are shown for HPA004754, HPA045140, HPA077723. (H+) refers to heat treated plasma and (H-)to untreated plasma . IGF2 was identified as P01344, and P01344-2 (*).

**Figure 4:**
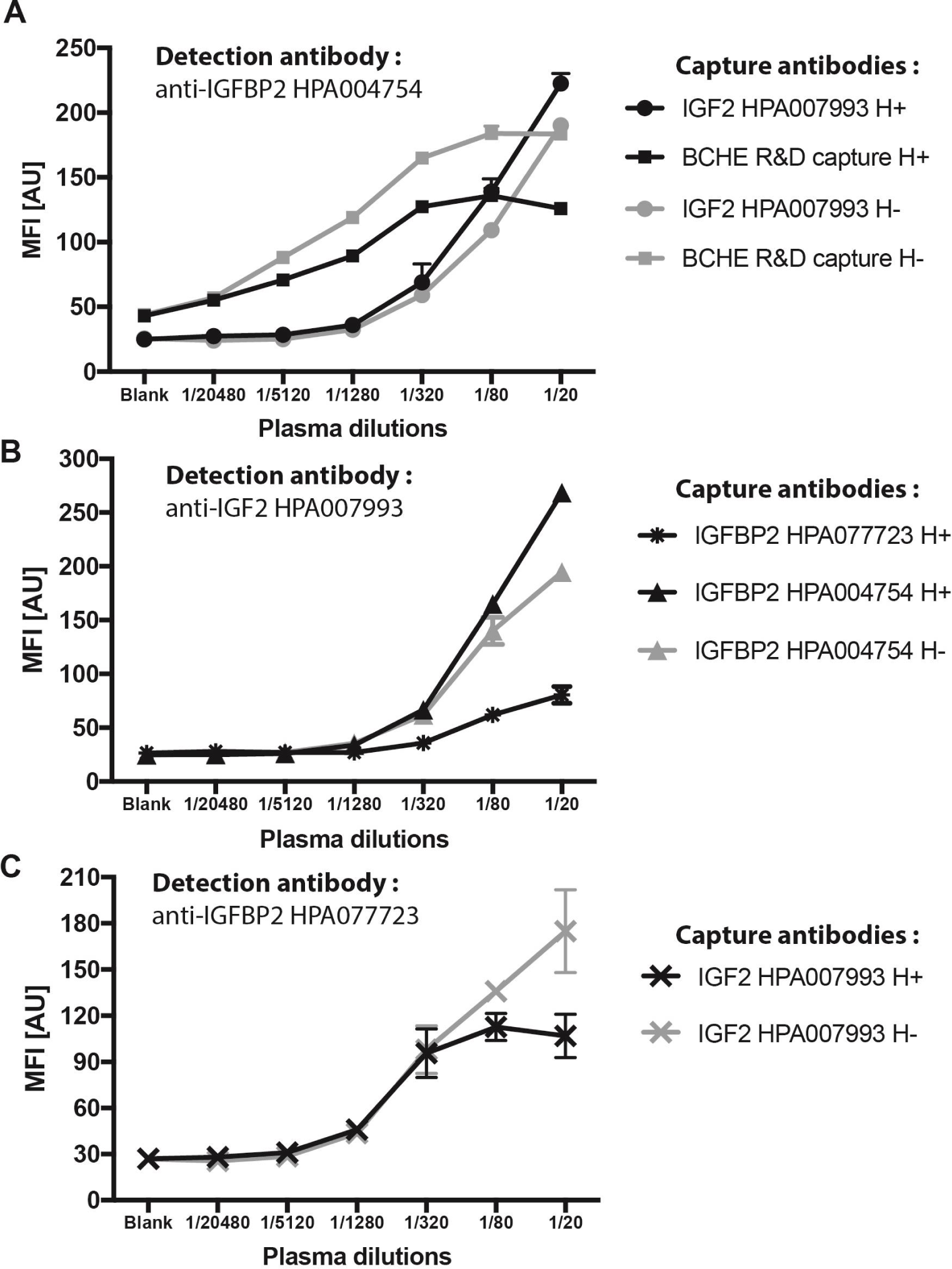
IGFBP2 protein interaction analysis by sandwich immunoassay. Dilution curves of plasma analyzed by sandwich assays using different combination of capture and detection antibodies. Dots represents mean value with standard error (SD) bars. In black, heated plasma (H+); in gray,non-heated plasma (H-).

Knowing that common plasma background proteins may differ between assays using untreated and heat-treated plasma, data from the respective assays was analyzed separately and the resulting z-scores were combined for the global assessment. For the 157 antibodies, we made 1173 identifications for 681 unique proteins with z ≥ 3. In each of the 414 IPs, we detected about 550 proteins, of which 6-9 proteins were enriched above the threshold. The combined outcome of the analyses is shown in **Table 1**, where the classification of 70 out of 157 antibodies (45%) was found supportive by falling into the ON-or CO-target category. When assessing only those antibodies targeting proteins previously annotated in plasma, the fraction increased to 61% (46/75). It is noteworthy that annotating antibodies in non-supportive categories could be due to limited affinity of the antibody, limited assay sensitivity or absence of the target in the used pools of plasma derived from healthy donors, as well as limited peptide detectability.

**Table 1:** Annotation and categorization and of antibodies.

| Annotated target location | On/Co-Target |  | Off/No-Target |  | Sum |  |
| --- | --- | --- | --- | --- | --- | --- |
|  | N | % | N | % | N | % |
| Cellular | 15 | 27 | 41 | 73 | 56 | 36 |
| Extracellular | 9 | 34 | 17 | 65 | 26 | 16 |
| Plasma (MS) | 46 | 61 | 29 | 39 | 75 | 48 |
| Total | 70 | 45 | 87 | 55 | 157 | 100 |

### ON-target enrichment

Applying IP-MS analysis has been reported to improve the sensitivity of protein quantification [8, 12, 25]. Hence, IP-MS may also be used to detect lower abundant proteins, adding to those that presently remain more challenging for other MS protocols. Our investigations lead to the identification of 9 extracellular proteins (e.g. CXCL8, TGFA) and 15 cellular proteins (e.g.TP53, CASP2) in plasma, that were not listed in the plasma PeptideAtlas at the time of our study **Supplementary Excel Sheet: “Antibodies experim. annotation”**. For almost 50 % of the polyclonal antibodies annotated as ON-or CO-target in our studies, the identified peptides aligned with the sequence of the protein fragments used to generate the antibodies or Protein Epitope Signature Tags (PrEST) (**Supplementary Fig.1B**). As previously discussed [26], affinity purification of polyclonal antibodies may bare the risk of coeluting the target used as bait on the columns and detecting these by MS. Such passenger proteins or peptides may consequently simulate the enrichment of an endogenous target. In our case, this would lead to a false classification of the antibody and may hamper downstream applications. To address this concern, we analyzed beads with immobilized antibodies for the presence of the protein fragment antigen that could cause passenger peptides to appear. Out of 42 tested antibodies (among ON-target or CO-target), only 11 showed the potential presence of passenger proteins (**Supplementary note 4, Supplementary Fig. 6**).

### CO-target enrichment and sub-categories

Evaluating antibodies in terms of target selectivity, we observed the possibility to study co-enrichments of proteins (**Figure 3**). This category includes those ON-target enrichments where other proteins were either enriched due to an interaction with the ON-target or as an OFF-target being captured by the antibodies alongside the indented target. For the second sub-category, sequence homology and abundance of the OFF-target could serve as reasons for co-enrichments.

### CO-target enrichment stemming from related proteins

Besides IGFBP2 discussed further below, examples of the CO-target enrichments with sequence homology included examples for anti-CCL16 (HPA042909) and anti-SERPINA4 (HPA002869). These binders respectively enriched additional members of their protein family, CCL18 and SERPINA6, both sharing sequence homology with the intended target (**Supplementary note 7**). The proteins CCL16 and CCL18 are estimated to be present at 29 ng/ml and 2 ng/ml levels in human plasma (Peptide Atlas, 2017), hence the estimated abundance differs from the degrees to which HPA042909 captured CCL16 (z = 10.7) and CCL18 (z = 10.1). Both proteins were otherwise rarely observed in other pIPs (*f* <4%), have not been predicted to interact directly (**Supplementary Fig. 7A**) but share sequence to a 27% homology (**Supplementary Fig. 7B**). In the case of HPA002869, the antibody enriched SERPINA4 (z = 8.2) and SERPINA6 (z = 8.1) in heated plasma. Serpins are a large family of blood proteins highly homolog proteins and no direct integration has been predicted (**Supplementary Fig. 7C**). SERPINA4 and A6 are estimated to be present at 17 and 41 μg/ml levels and share a 40% sequence similarity (**Supplementary Fig. 7D**). Both were observed in 91% (SERPINA4) and 59% (SERPINA6) of all conducted pIPs with heat-treated plasma (less frequent in untreated plasma). A research performed on PIPs, a database of predicted human protein-protein interactions (http://www.compbio.dundee.ac.uk/www-pips/index.jsp, [27]), revealed a predicted interaction between SERPINA4 and SERPINA6 (total interaction score = 128.0). Further antibodies with independent epitopes are needed to confirm or to exclude the possibility of interactions between these proteins, as we did not find literature supporting a physical interaction between them.

### CO-target enrichment stemming from frequent proteins

In another sub-category, we observed that potential protein-protein interactions may also be described from more frequently observed proteins that tend to present with z-scores <3. Below, we discuss two examples found for IGM, a protein known to be abundant in plasma. The first example is given by CD5 antigen-like (CD5L, HPA026432) and its known interactor IGM. Here, CD5L was detected alongside several immunoglobulins (IGH, IGJ, IGK, IGL, IGM). It is known that CD5L takes part in inflammatory responses during infections or during the process of atherosclerosis, hence binds to the Fc region of IgM through its SRCR domains [30]. Further, the immunoglobulin J chain (IGJ) is known to be required to stabilize the binding of CD5L to IGM, but a direct interaction has not been experimentally observed [31]. Our data supports the idea that association between CD5L, IGM and IGJ can occur: Besides CD5L ([c] = 5.9 μg/ml; z = 5.1), immunoglobulin light chain lambda (IGHV3-23; z = 3.4), kappa (IGKC; z = 3.4), IGM (z = 2.7) and IGJ (z = 2.8). A lower z-score of the co-targets may also possibly indicate that the antibody is more selective for CD5L and less for the additionally identified proteins. Considering the abundance of IGM at around 1 mg/ml and that IGM frequently occurred as contaminant (*f* = 98%), an increased z-score of IGM in this particular IP may point to a specific enrichment. The second example is given for a monoclonal antibody raised against fibulin 1 (FBLN1) [32]. This antibody was used in both sample types and enriched 22 proteins in heated and 12 in untreated plasma. Even though FBLN1 is a frequent protein (*f* = 65%) and abundant protein ([c] = 34 μg/ml), it was only enriched in heated plasma (z = 3.7), while other targets such as CAPZA2, IL36G and PLK4I were detected in both preparations (**Supplementary Excel Sheet: “Antibodies against same protein”**). Interestingly, CD5L and IGHM were again found in the same assays, with comparable enrichment values in heated (z = 3.4 and 3.3) compared to untreated plasma (both z = 2.9). This could serve as another indication that CD5L and IGHM are present in a complex in plasma. Further analyses are though needed to determine the mode of co-enrichment, meaning, if the co-target was enriched due to directly interacting with the on-target or due to be serving as an off-target.

### Studying protein interaction with paired antibodies

The biologically most interesting sub-category within the CO-target category refers to proteins that co-identified because they presumably interact with the intended target. We chose a limit in z ≥ 3 and LFQ intensity ≥ 10^7^ to call potential protein interactions. In general, we observed that most of the consistent identifications (identified in several replicates) were found for LFQ precursor intensities above this level (**Supplementary Fig. 10**). One such example is given by the pIP using three antibodies raised against the insulin growth factor binding protein 2 (IGFBP2: HPA077723, HPA045140, HPA004754) of which the latter two were raised against the same antigen. As shown in **Fig. 3B** for untreated plasma, HPA077723 and HPA045140 both enriched IGFBP2 ([c] = 1.1 μg/ml; *f*=21%) as well as previously known interactors insulin growth factor 1 (IGF1: [c] = 0.46 μg/ml; / = 18%;) and IGF2 ([c] =1.6 μg/ml; 8%). In addition, the plasma proteins butyrylcholine esterase (BCHE: [c] = 11.0 μg/ml; *f* = 18%;) and the deoxyribose-phosphate aldolase (DERA: [c] = 0.5 ng/ml; *f* = 7%) were detected. For BCHE and IGF1, an interaction was indeed previously hypothesized [28, 29]. For the third binder (HPA004754), IFGBP2 was only enriched upon prior heat treatment plasma (**Fig. 3B**). This differential performance of antibodies raised against the same antigen (HPA045140, HPA004754) confirms the necessity to investigate each of the different batches and lots of polyclonal antibodies.

In order to provide further evidence for an interaction between the identified proteins, we conducted multiplexed sandwich assays. Here, recombinant IGFBP2, IFG1, IGF2 and BCHE were first analyzed in a concentration dependent manner to confirm assay functionality and target specificity (**Table 2**). First, EDTA plasma was analyzed in a concentration dependent manner, confirming the selectivity of the matched antibody pairs (**Supplementary Fig. 11**). Then, we investigated if antibody pairs with different selectivity revealed plasma concentration dependent results. As shown in **Fig 4**, we found pairs of antibodies mixed specificity in the following capture-detection configurations: IGFBP2-IGF2, IGF2-IGFBP2 as well as BCHE-IGFBP2. For IGF1 and IGF2 antibody pairs, it was not possible to obtain a dilution curve with the respective recombinant proteins in solution, but they were functional in plasma (**Supplementary Fig. 11 C,D,M,N,and P**)). Also, IGF2-IGFBP2 and IGFBP2-IGF1 confirmed the presence of the previously known complex IGFBP2-IGF2 (**Table 2, Fig 4 B-C**). Antibody pairs for IGFBP2 and BCHE described a sample dilution depended trend with their corresponding intended recombinant proteins as well as in plasma (**Table 2 and Supplementary Fig. 11A, E,F,H and I**)). Since no cross-reactivity was observed towards these two proteins with other antibodies in the assay (**Table 2**), the functional antibody pair BCHE-IGFBP2 supports the indications provided by IP that a physical interaction between the two proteins in plasma (**Fig 4A**). An inverted configuration IGFBP2-BCHEand an assay including also the other IGFBP2 antibody indicating an interaction (HPA077723, **Fig. 3B**) would further support this observation. However, we acknowledge that not all antibodies allowed building mixed sandwich pairs and using the chosen assay protocol, and indeed, HPA004754 and HPA077723 were raised against two different regions of IGFBP2. This could explain their different performance as capture and detection, above all in the presence of complexes containing IGFBP2. HPA004754 was though functional as capture and detection antibody both using heat-treated and untreated plasma for the detection of IGFBP2 and in combination with anti-BCHE (**Supplementary Fig. 11. B and G**). HPA077723 was though not functional with anti-BCHE either as a capture or detection antibody, suggesting that the binding of either antibody hinders the other antibody binding to IGFBP2-BCHE complex. Further investigations are needed to investigate if this hindrance is due to a proximity of the two binding sites or other steric effects such as epitope accessibility of a captured complex.

**Table 2:**
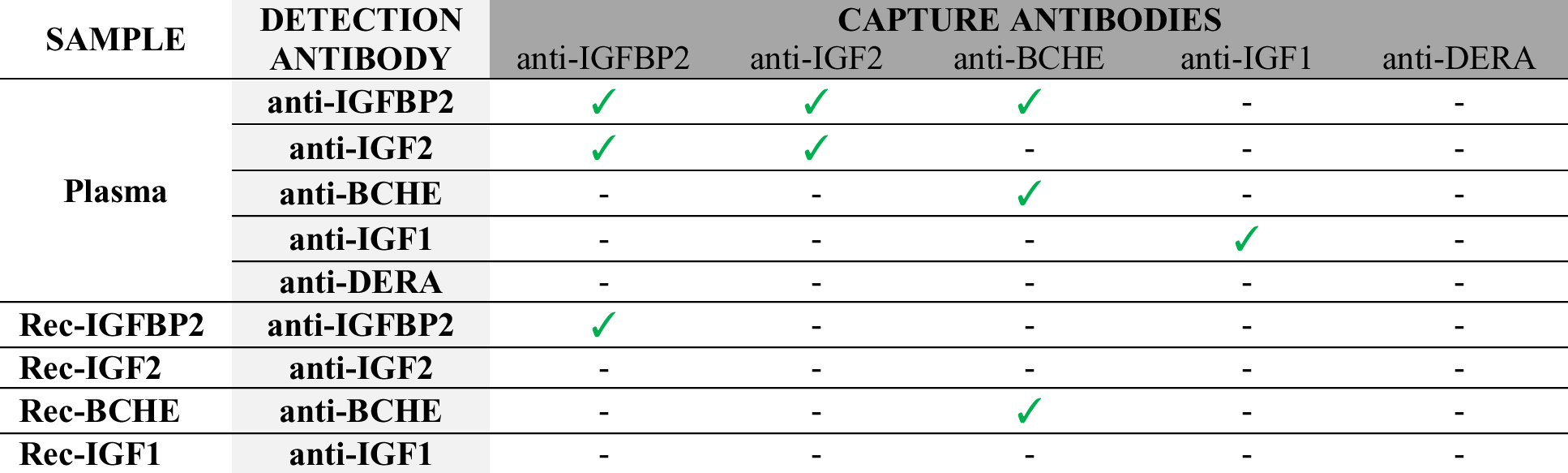
Antibody pairs tested in plasma and with recombinant proteins. Annotation (✓): Trends from sample dilution assays were obtained with at least one combination of antibodies for the same protein either in heated or untreated samples. Annotation (-): No sample concentration dependent data was obtained. (Details regarding each single pair in Supplementary Excel Sheet “Reagents_Lot_Numbers”, catalog numbers of the functional pairs are indicated in **Figure 4** and **Supplementary Figure 10**.

More examples of potential protein interactions are provided by those cases in which multiple antibodies were raised against the same target protein using different antigens. Enriching common ON-target and CO-targets provides evidence for protein complexes rather than artefacts. Examples are antibodies targeting FBLN1, or IGF1R (see **Supplementary Excel Sheet: “Antibodies against same protein”**).

### OFF-target enrichment category

As a last category, we investigate off-target enrichments. Here, we see the plasma abundance of the off-target over the intended analyte as a main reason for failing to enrich the expected target. As the community is starting to acknowledge the fact that the performance of antibodies is indeed sample context and application dependent, certifying off-target interactions may still allow generating novel hypotheses given that these are followed-up and thoroughly validated by appropriate targeted analysis. One example for selective off-target interactions in plasma is presented by the antibody HPA004920, raised against MMP1 (Matrix metalloproteinase 1). We classified this antibody as OFF-target in untreated plasma because it enriched Mannose-binding protein C (MBL2; z = 8.3; *f*=12%) as well as MMP3 (z = 6.8; *f*=6%) in the IP assays (**Figure 2C**). As described above for CCL16 and SERPINA4, also here a 53% sequences similarity between the intended target (MMP1) and the off-target (MMP3) (**Supplementary Fig. 7F**) exists and an interaction between these two proteins has been predicted (**Supplementary Fig. 7E**). The other off-target MBL2 and MMP1 have only a 10% sequence similarity (**Supplementary Fig. 7G**). MBL2 is though estimated to be present at 1.7 μg/ml in circulation, it is almost 1000x more abundant than MMP1 ([c] = 1.1 ng/ml) and MMP3 ([c] = 0.5 ng/ml) [14]. MBL2 has also been described in to reside in a complex with the serine protease MASP (MBL-associated serine protease) [31], and its collagen-like domain may serve as a substrate for matrix metalloproteinases (MMPs) to nest in. Studies on MBL mutations suggest that MMPs may be involved in physiological regulation of MBL levels [32]. This could eventually explain the presence of MBL2-MMP3 complexes in plasma.

### Conclusion

In summary, this study describes a resource that was built from our interest in the verification of antibodies selectivity in plasma. The antibodies analyzed in this study include polyclonal and monoclonal that was used for exploratory bead arrays and the development of immunoassays, where either heat-treated or untreated plasma may serve as samples. We have conducted > 400 IP assays in plasma and built a library of proteins with their frequencies of identification in plasma. Constructed on the systematic analysis of 157 antibodies, we described the occurrence of common proteins, denoted plasma background, which allowed us to determine the selectively captured endogenous plasma proteins by mean of z-scores analysis. Our approach, which we also compared with Western blot (**Supplementary note 5** and **Supplementary Table 1**), could serve as a valuable method to narrow large numbers of antibodies determining which ones could enrich the endogenous protein of interest and to further investigate assay selectivity as well as proteins interactions.

This concept may though not yet elucidate (i) if the antibodies bind to proteins at full-length or fragments, (ii) if the antibodies will be functional in pairs in sandwich assay, (iii) how potential protein interactors and off-targets would compete with on-target binding, (iv) how contaminations from passenger antigens affect the assay’s selectivity and sensitivity. For ON-target and CO-target antibodies, thus further investigations are required to clarify the technical aspects mentioned above and to expand on the biological implication of protein interactions in plasma. Nevertheless, pIP is an informative first test for the identification of pair antibodies with different target proteins to study protein complexes in plasma by sandwich assays. While, targeted fit-for-purpose experiment should then include dose response curve in dilutions of plasma, preferentially coupled to quantitative mass spectrometry analysis for a set of identified peptides including potential co-targets and/or contaminants.

The provided resource builds one foundation towards a more detailed assessment of antibodies for plasma proteomics assays, and may contribute to the development and application of more specific, robust and reliable immunoassays that can use mass spectrometry or other means of detection [1].

## MATERIAL AND METHODS

Methods are available in the online version of the paper.

## AUTHOR CONTRIBUTIONS

CF and JMS initiated, designed, and led the study with the scientific contribution and support from all co-authors. CF designed the IP experiments and performed the data analysis. CF, SB, LSR and MI executed the IP experiments with support from DT and RMB.LSR conducted the immunoassays. JL provided MS infrastructure. The study was supervised by PN, JL and JMS. CF and JMS wrote the manuscript with input from all authors.

## AKNOWLEDGMENTS

We greatly thank Mathias Uhlén, Fredrik Edfors, Björn Forström, Lucia Lourido, Gabriella Tekin and the Biobank profiling, Affinity Proteomics and Clinically Applied Proteomics groups at SciLifeLab in Stockholm for their continuous fruitful discussion, access to instrumentation and input to the presented work. We also thank everyone at the Human Protein Atlas for their support, and thank Hanna Tegel, Johan Rockberg and their team for recombinant proteins. The KTH Center for Applied Precision Medicine (KCAP) funded by the Erling-Persson Family Foundation is acknowledged for financial support. This work was supported by grants for Science for Life Laboratory, the Knut and Alice Wallenberg Foundation. The work leading to this publication has received support from the Innovative Medicines Initiative Joint under grant agreement n°115317 (DIRECT), resources of which are composed of financial contribution from the European Union's Seventh Framework Programme (FP7/2007-2013) and EFPIA companies’ in-kind contribution. Funding from the Swedish Foundation for Strategic Research, Swedish Cancer Society and Swedish Research Council is gratefully acknowledged.

## ABBREVIATIONS

HPA: Human Protein Atlas
LFQ: label free quantification
PrEST: Protein Epitope Signature Tags
Z: z-score
*f*: frequency of occurrence

## SUPPLEMENTARY FIGURES & TABLES

**Supplementary Figure 1A:**
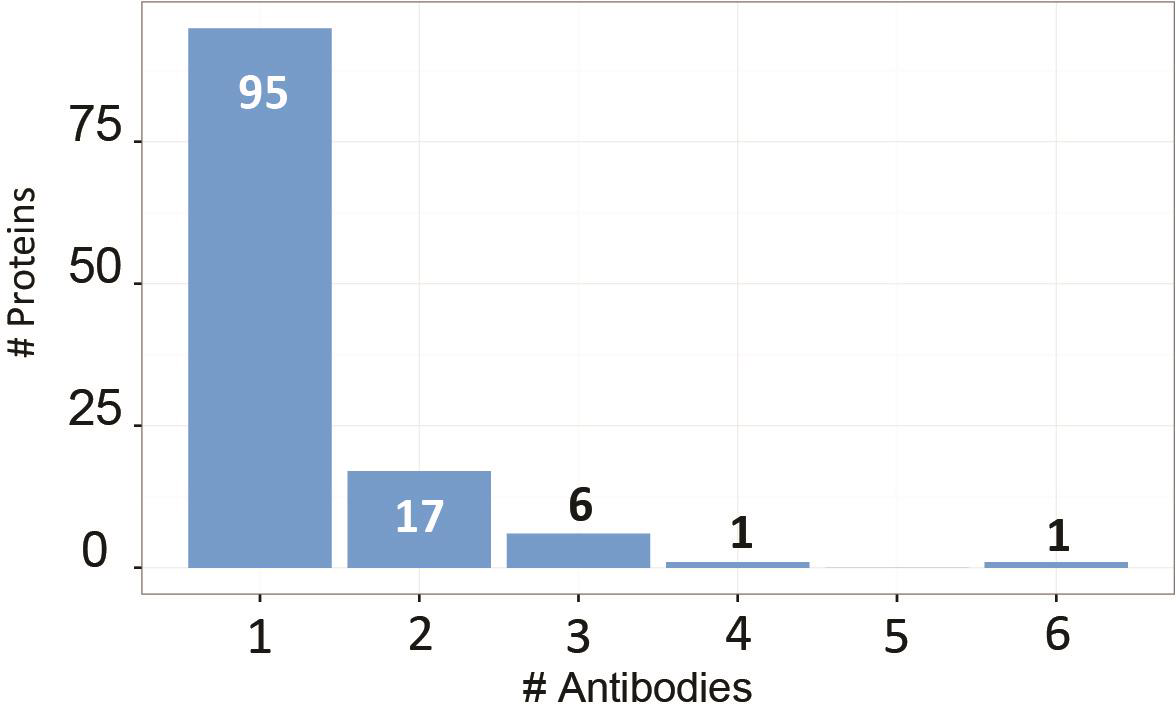
Distribution of number of antibodies targeting each protein.

**Supplementary Figure 1B:**
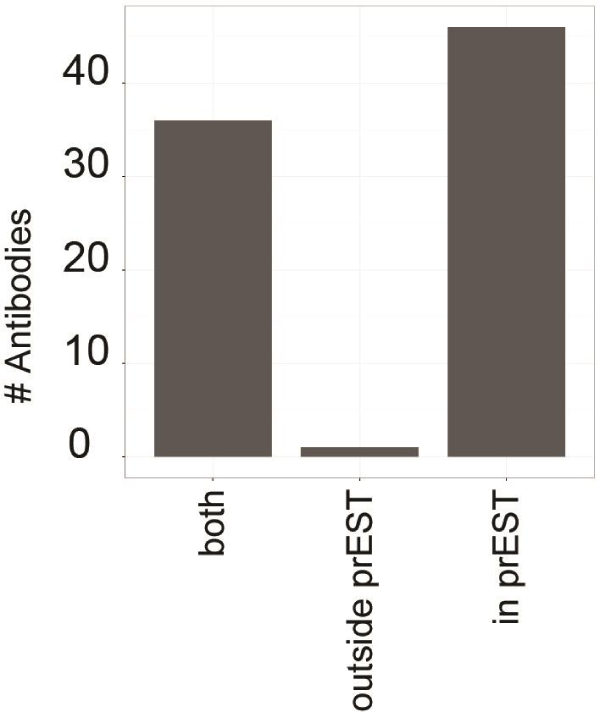
Coverage of proteins identified. Distribution of ON-target and CO-target antibodies based on the peptides identified for the intended target: peptides only covering the PrEST sequence (in PrEST), peptides not covering the PrEST (outside PrEST) or peptides identified inside and outside PrEST sequence (both).

**Supplementary Figure 2A:**
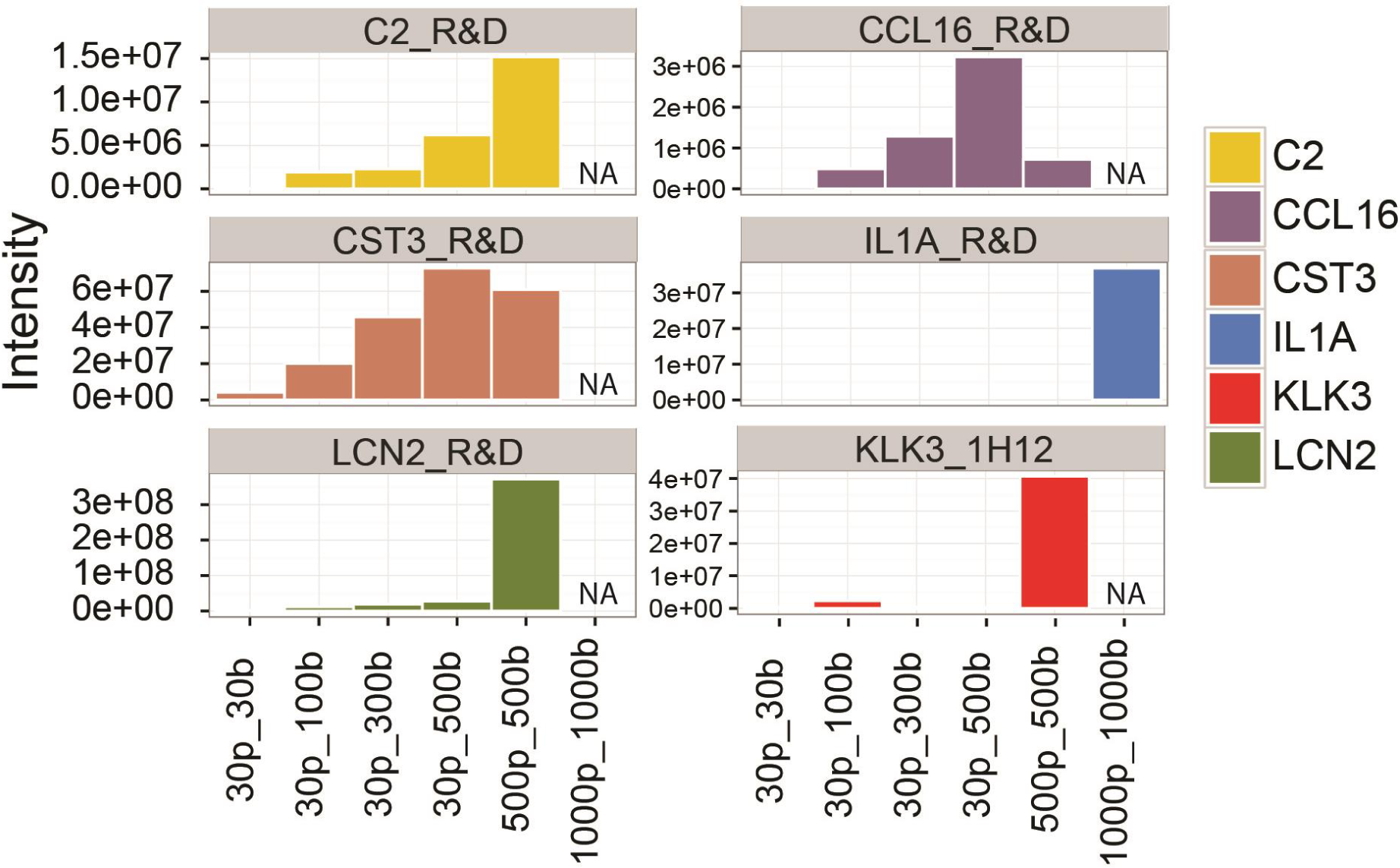
Optimization of experimental and technical conditions. Commercial antibodies targeting known plasma proteins of different abundance (C2, CCL16, CST3, IL1A, KLK3, LCN2) were used to establish optimal experimental conditions for volume of plasma and number of beads coupled to antibody. “p”=volume of plasma in μL; “b”= number of coupled beads (500000b = 1.6 μg of antibody).

**Supplementary Figure 2B:**
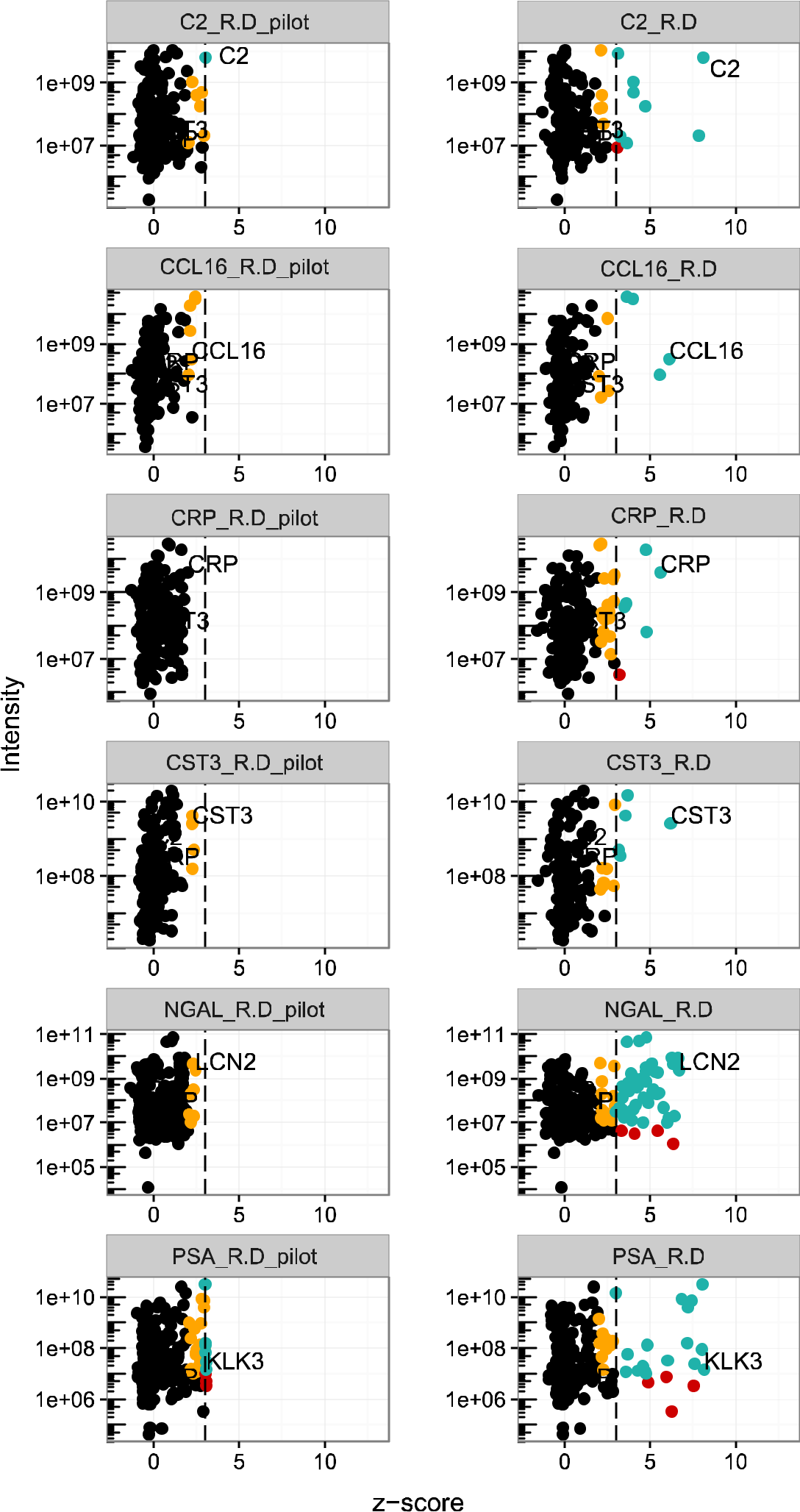
Effect of library size. Data derived from IP assays with antibodies targeting the plasma proteins of C2, CCL16, CRP, CST3, NGAL and KLK3 (PSA) was analyzed using the pilot library limited to only these 21 IPs (left column,'’’pilot”) as well as the library of more than 400 IP (right). In this experiment, 100 μl of plasma and 500,000 beads were applied.

**Supplementary Figure 3 A:**
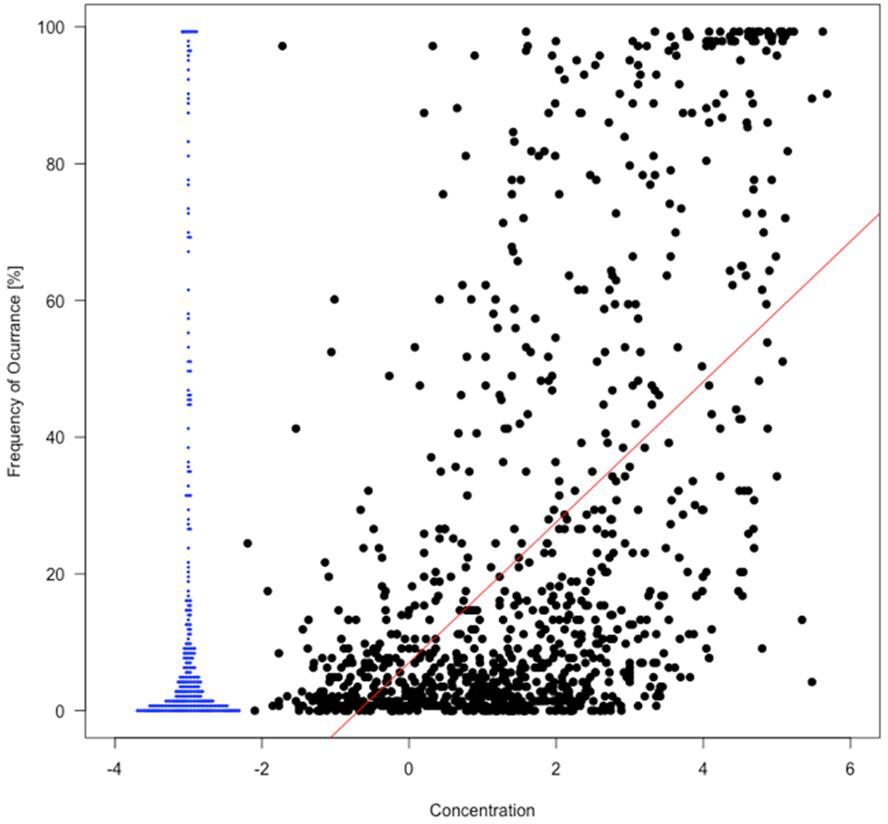
Relation between estimated plasma concentration [ng/ml] and frequency [%] for untreated plasma. Blue dots represent proteins for which estimated values of concentration were not present in PeptideAtlas.

**Supplementary Figure 3 B:**
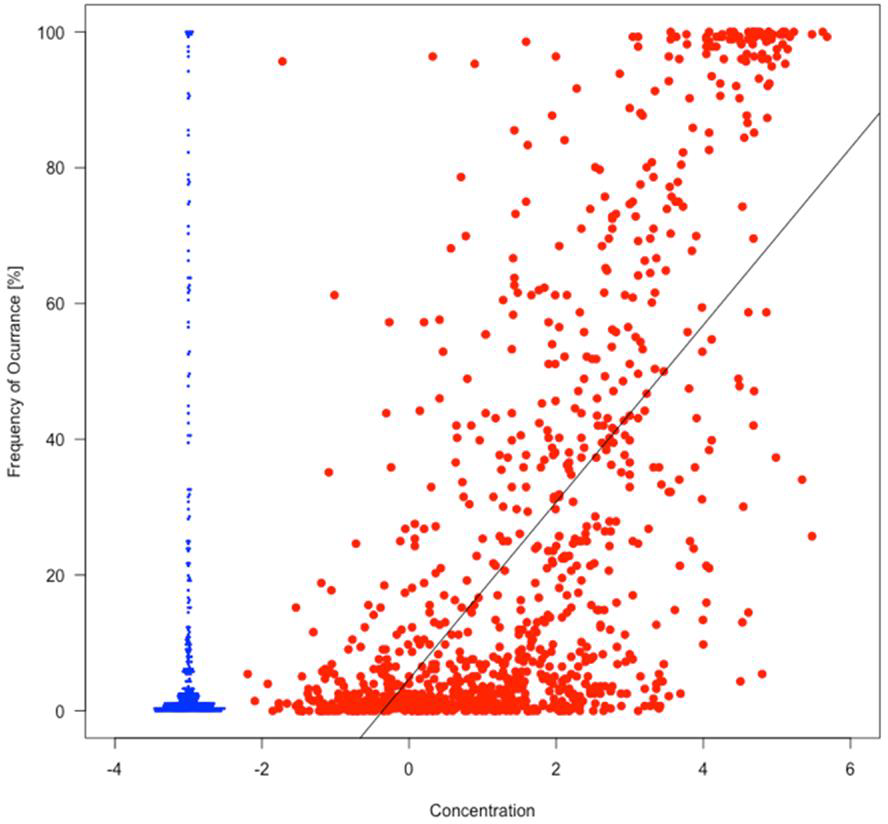
Relation between estimated plasma concentration [ng/ml] and frequency [%] for heat-treated plasma. Blue dots represent proteins for which estimated values of concentration were not present in PeptideAtlas.

**Supplementary Figure 4 Evaluation of contaminant proteins identified by pIPs and comparison of experimental batches.**

**Supplementary Figure 4 A:**
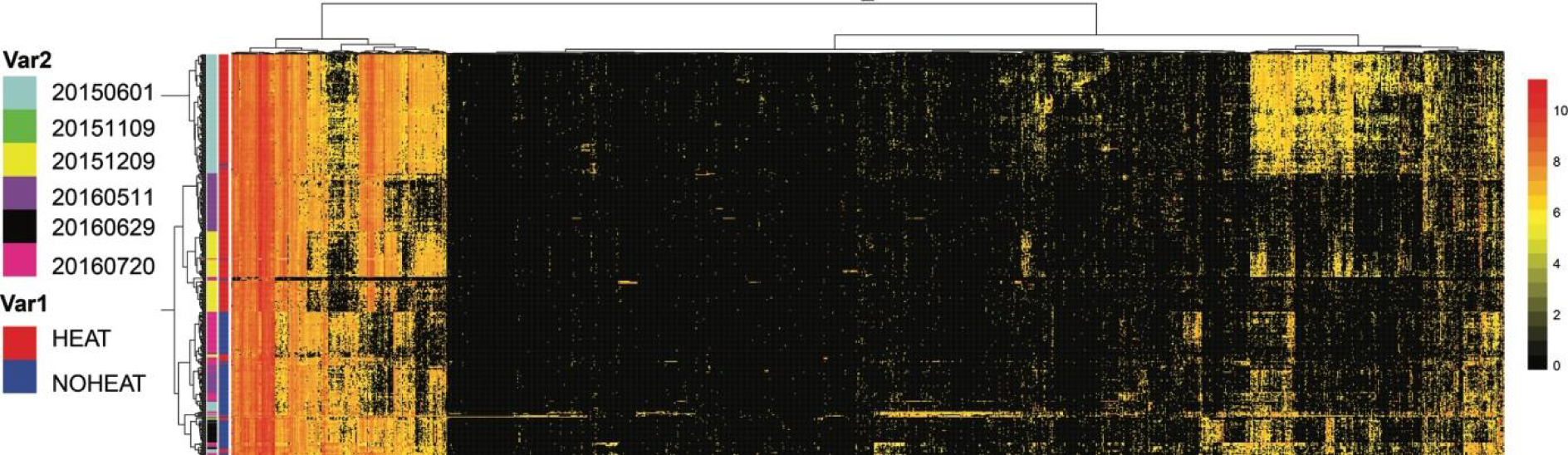
Two ways hierarchical clustering analysis of the proteins identified in 414 immunoprecipitations. Clustering distance: "euclidean”; Clustering method: Ward. Cluster bars represent (1) different experimental batches and (2) sample heat treatment.

**Supplementary Figure 4 B:**
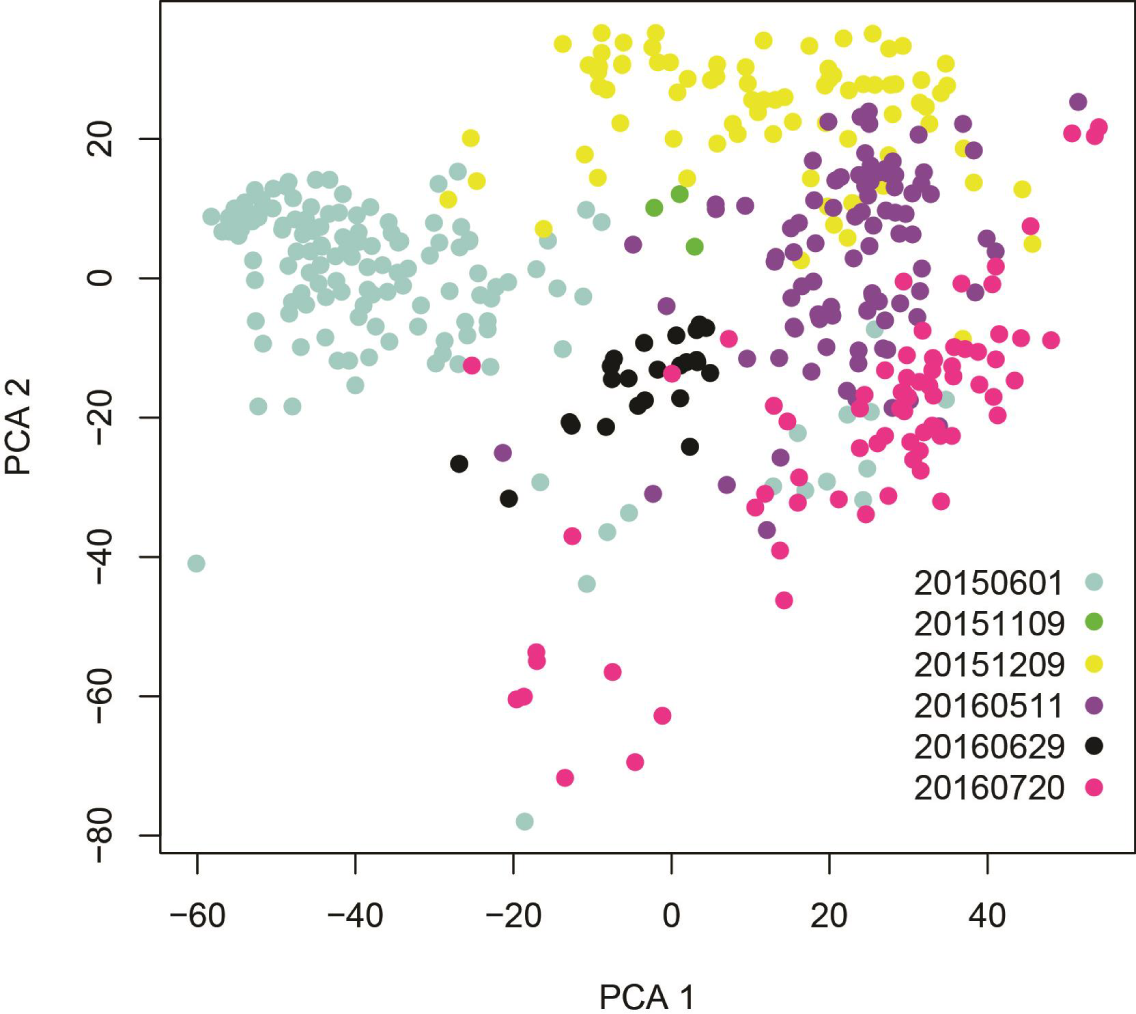
Representation of principal component analysis (PCA) with dots colors indicating the experimental batches.

**Supplementary Figure 5.**
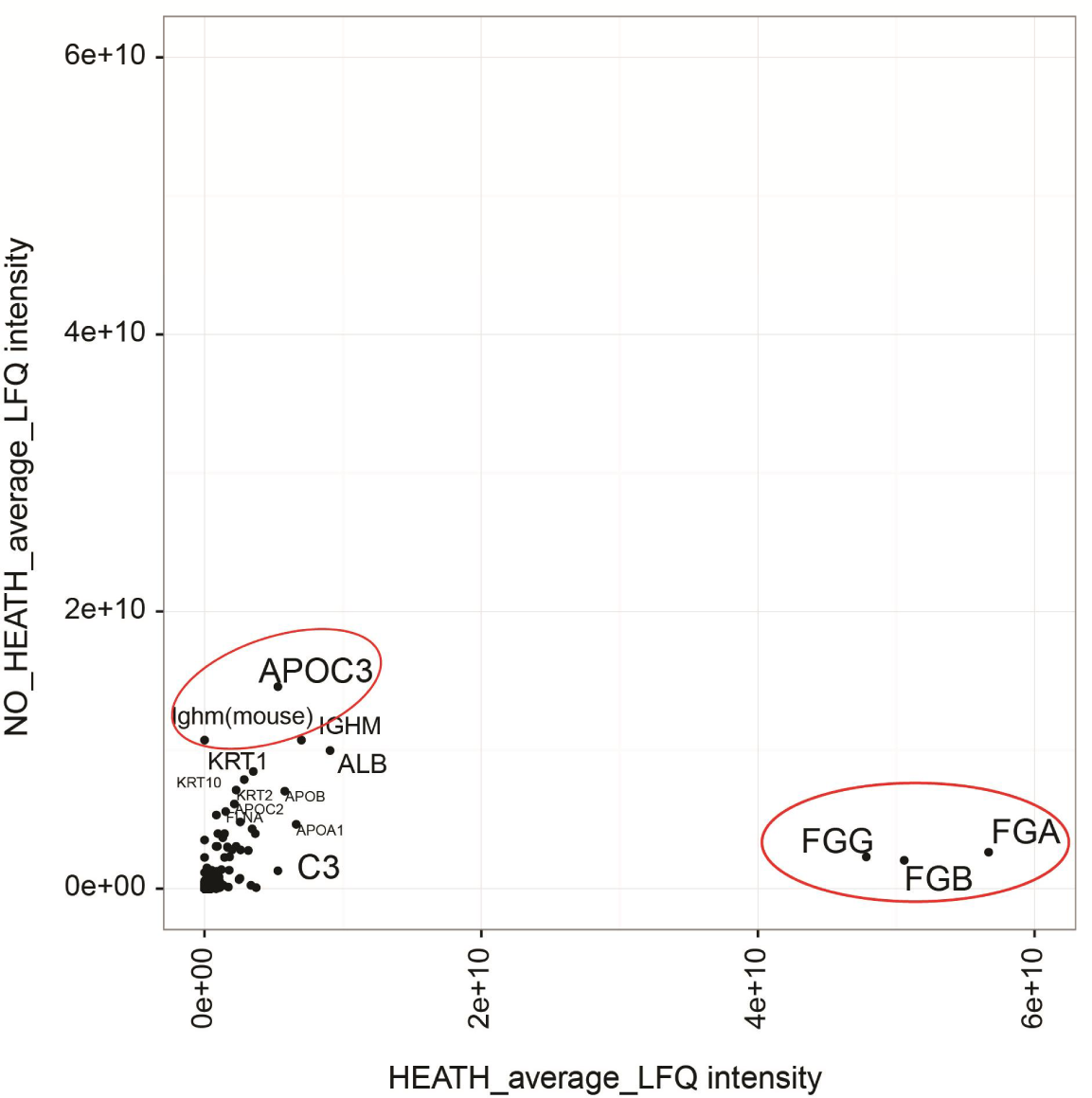
Comparison of LFQ protein intensities for IMS performed in heat treated versus in not heat treated plasma. Circled in red high abundant plasma proteins which abundance in the background increases when plasma is heat treated.

**Supplementary Figure 6:**
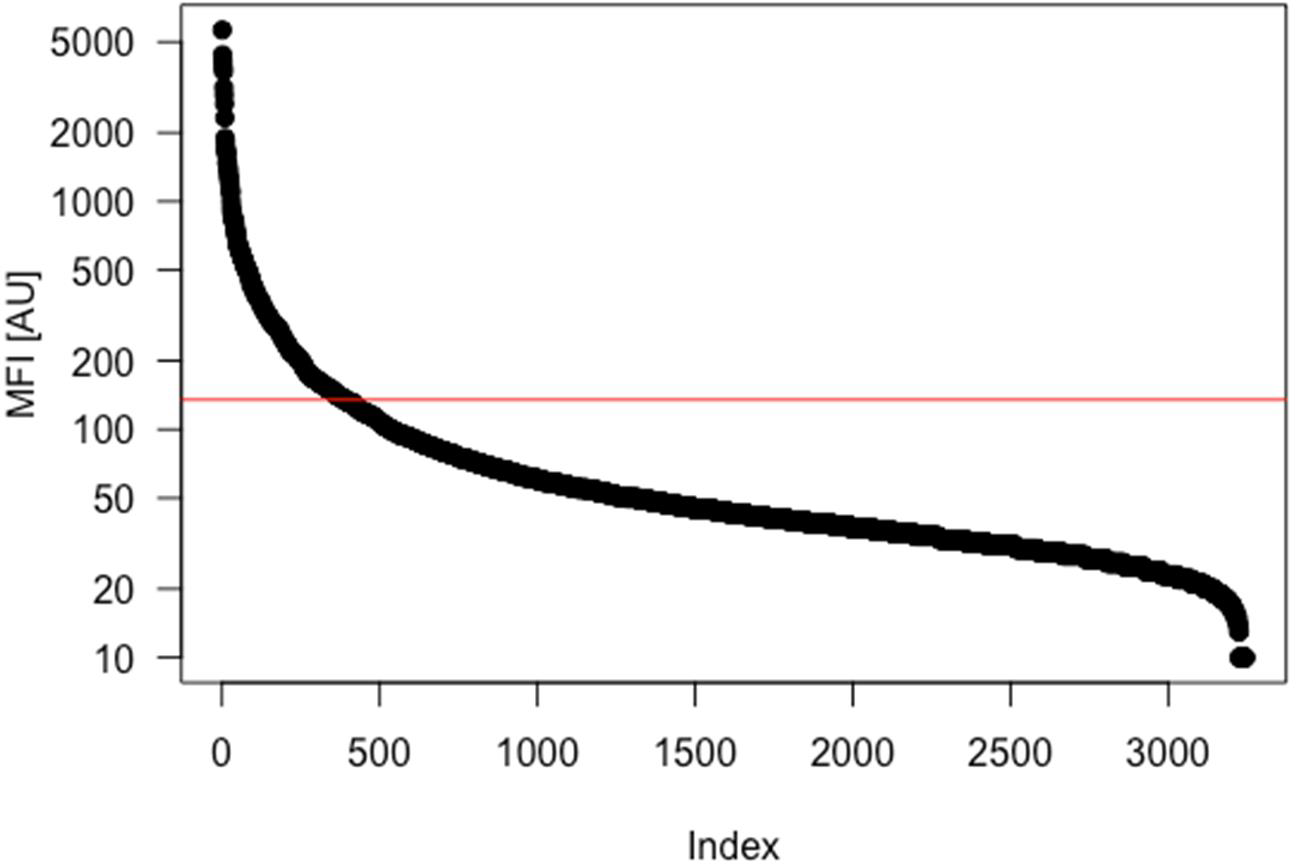
Ranked distribution of the detection of passenger antigens on antibody coupled beads.

**Supplementary Figure 7A:**
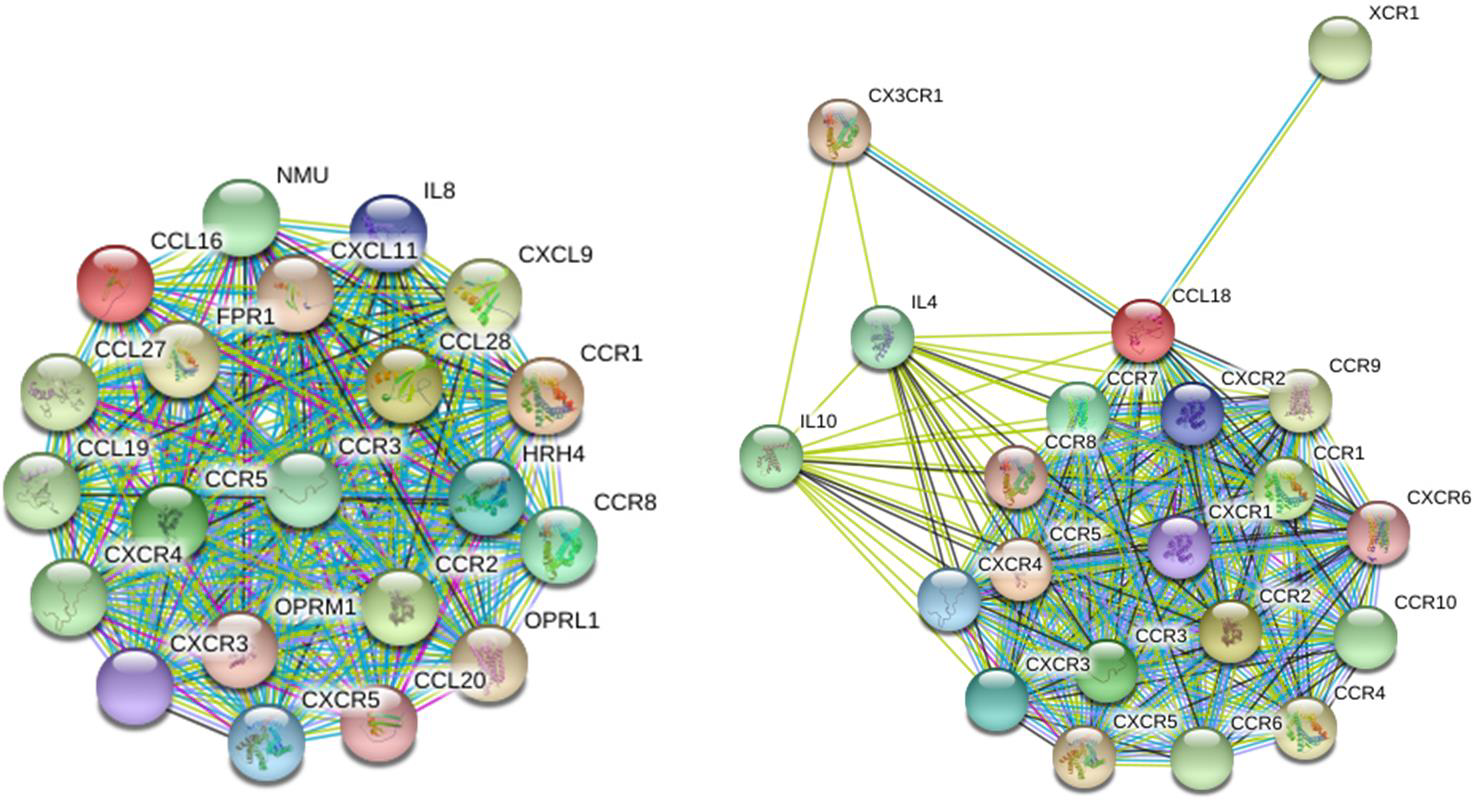
Interaction networks of CCL16 and CCL18 taken from String database V10.5 (20 interactors, 1^st^ shell only).

**Supplementary Figure 7B:**
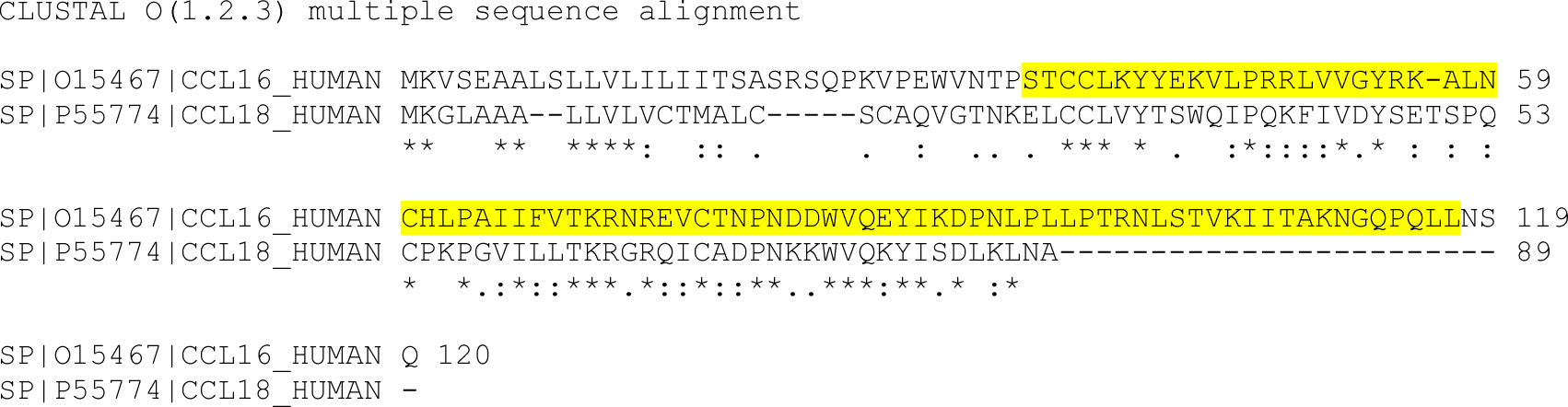
Sequence homology search of CCL16-CCL18 using CLUSTALO alignment revealed a 26.6% homology. In Yellow: Amino acidic sequence used as antigen in the generation of the antibody (PrEST).

**Supplementary Figure 7C:**
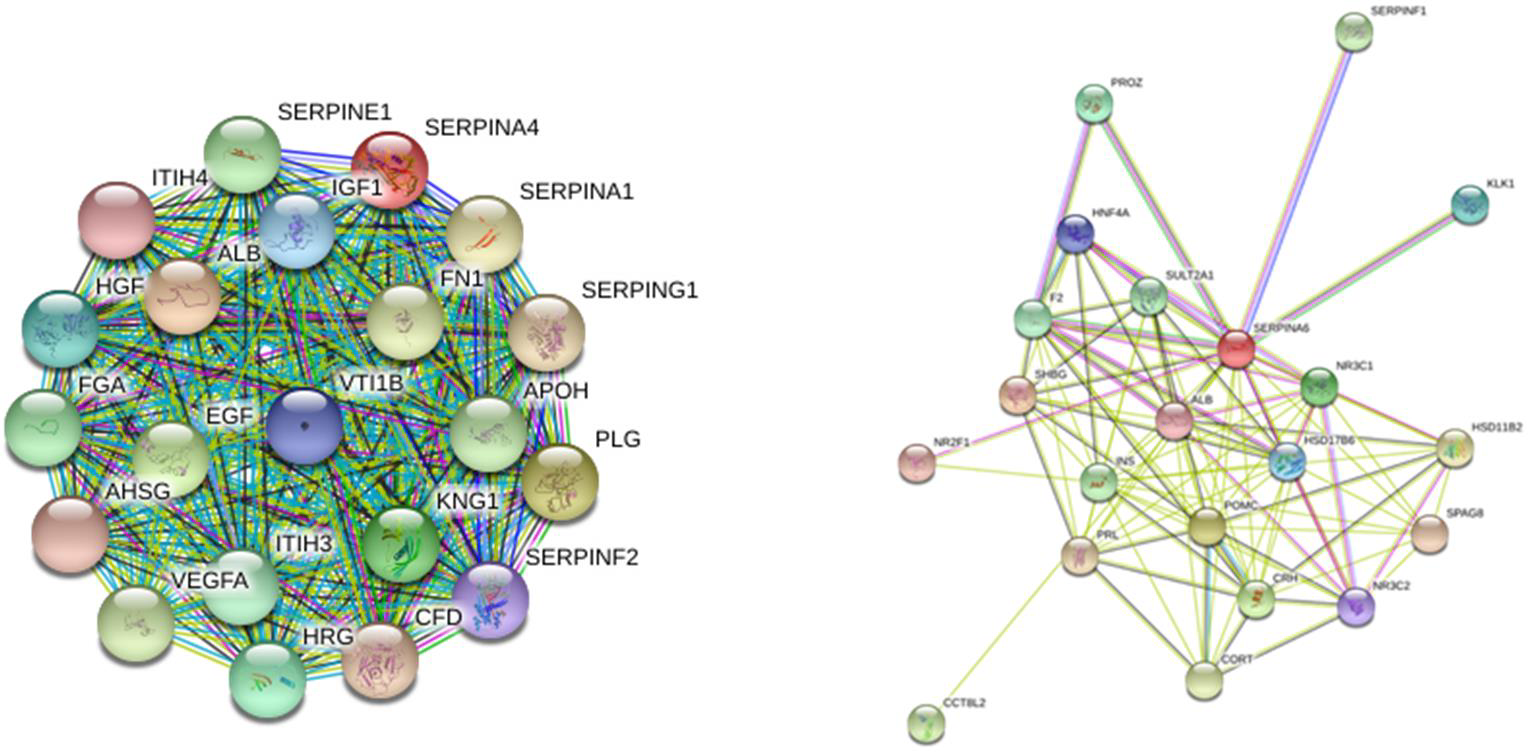
Interaction networks of SERPINA4 and SERPINA6 taken from String database V10.5 (20 interactors, 1^st^ shell only).

**Supplementary Figure 7D:**
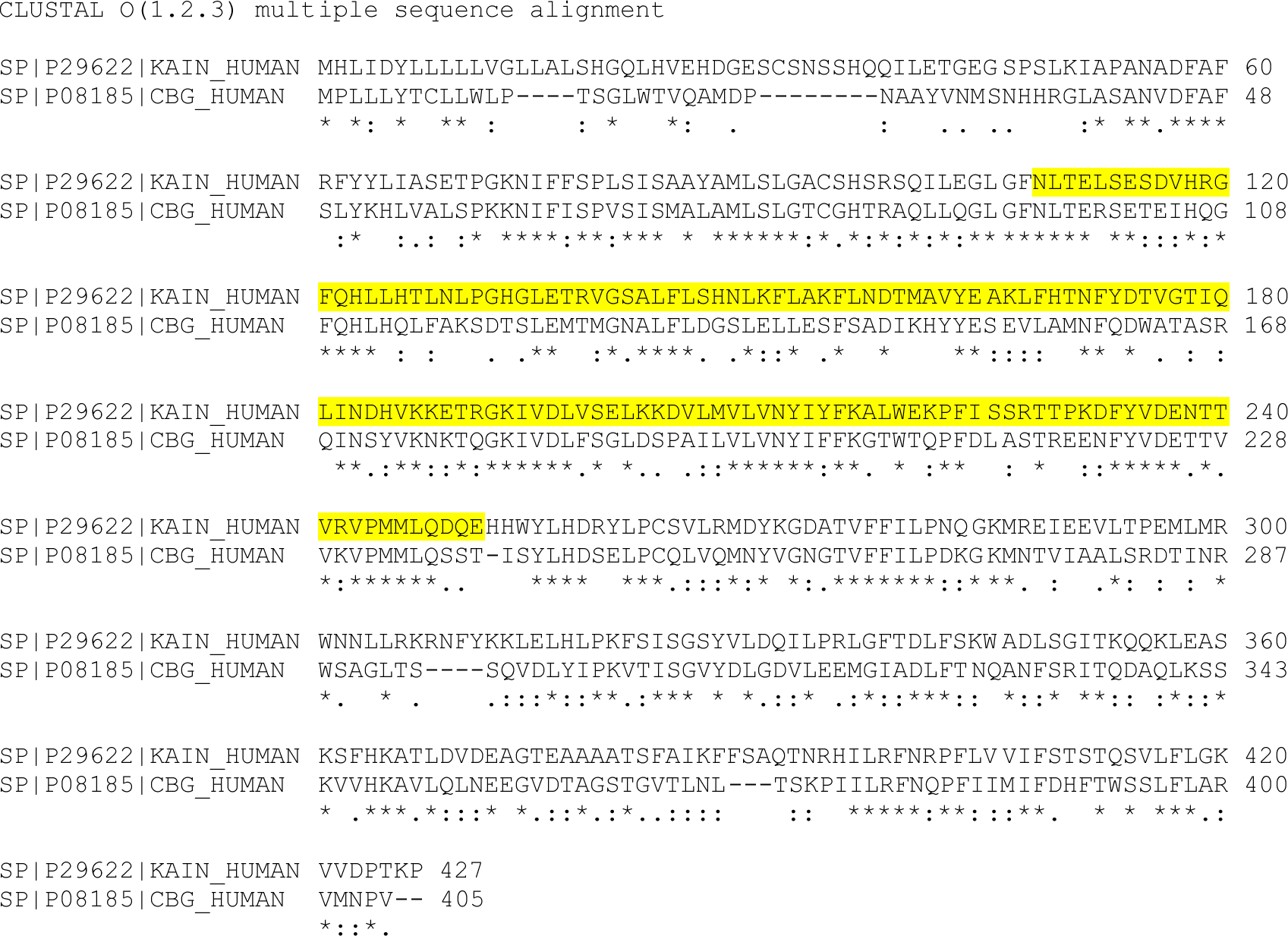
Sequence homology search of SERPINA4-SERPINA6 using CLUSTALO alignment revealed a 40.0% homology. In Yellow: Amino acidic sequence used as antigen in the generation of the antibody (PrEST).

**Supplementary Figure 7E:**
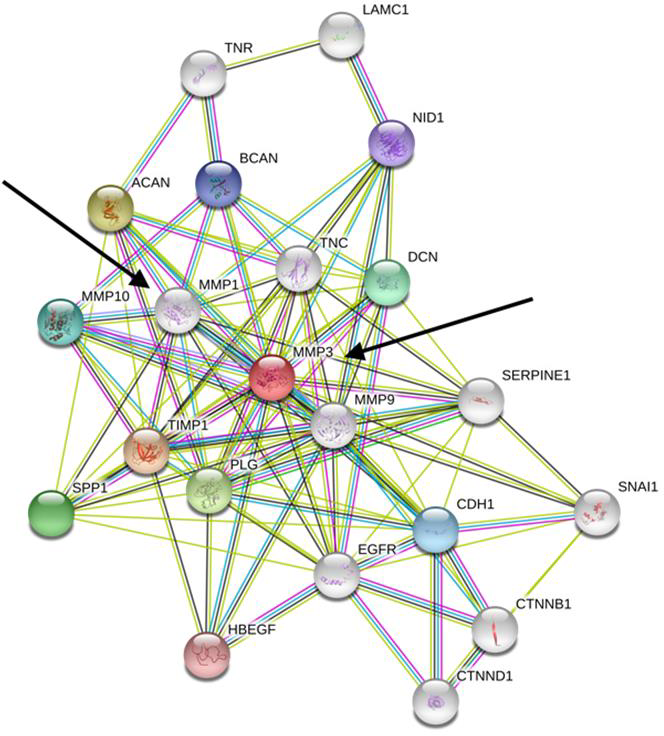
Interaction network of MMP1 and MMP3 taken from String database V10.5 (20 interactors, 1^st^ shell only).

**Supplementary Figure 7F:**
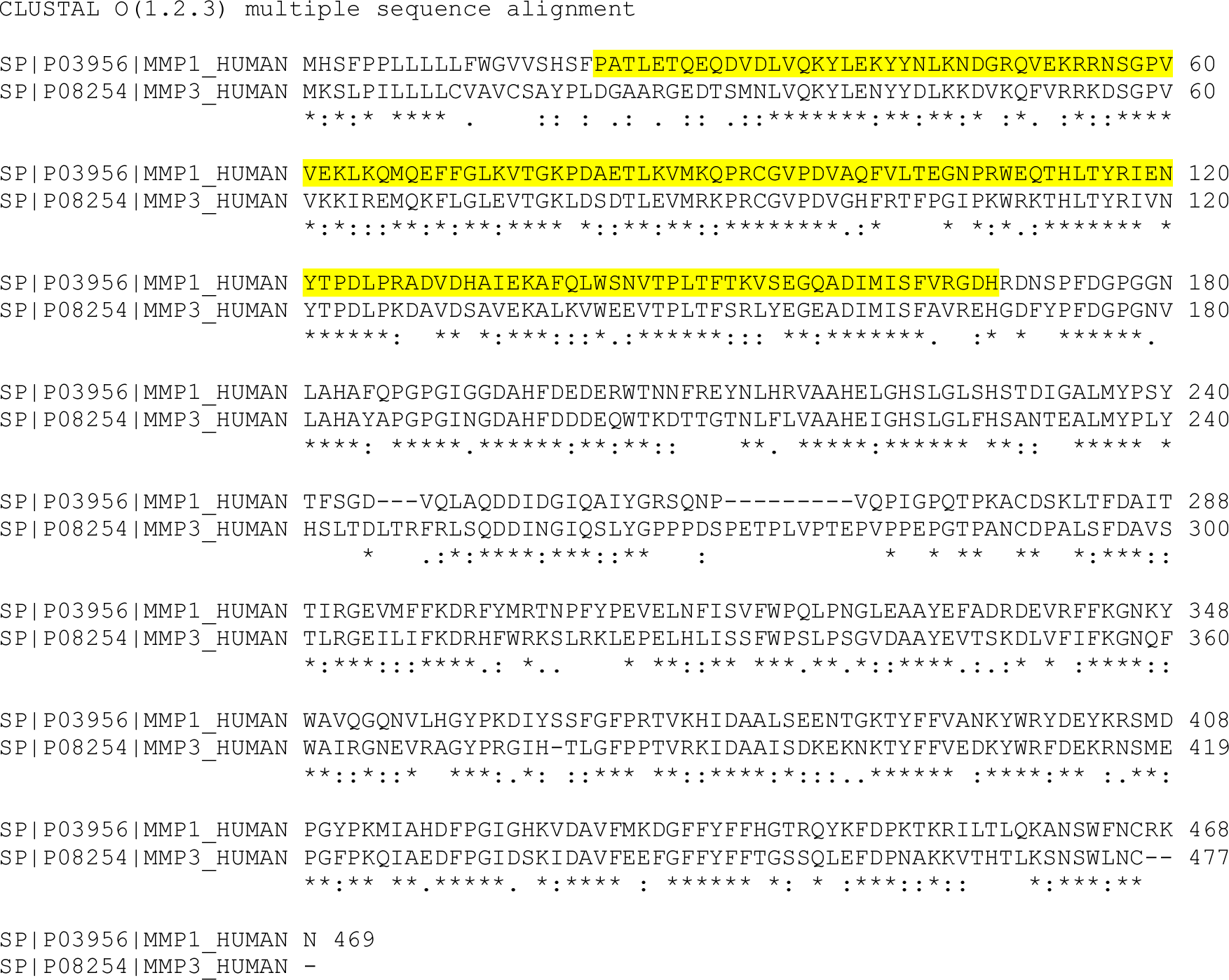
Sequence homology search of MMP1-MMP3 using CLUSTALO alignment revealed a 52.8 % homology. CLUSTAL. In Yellow: Amino acidic sequence used as antigen in the generation of the antibody(PrEST).

**Supplementary Figure 7G:**
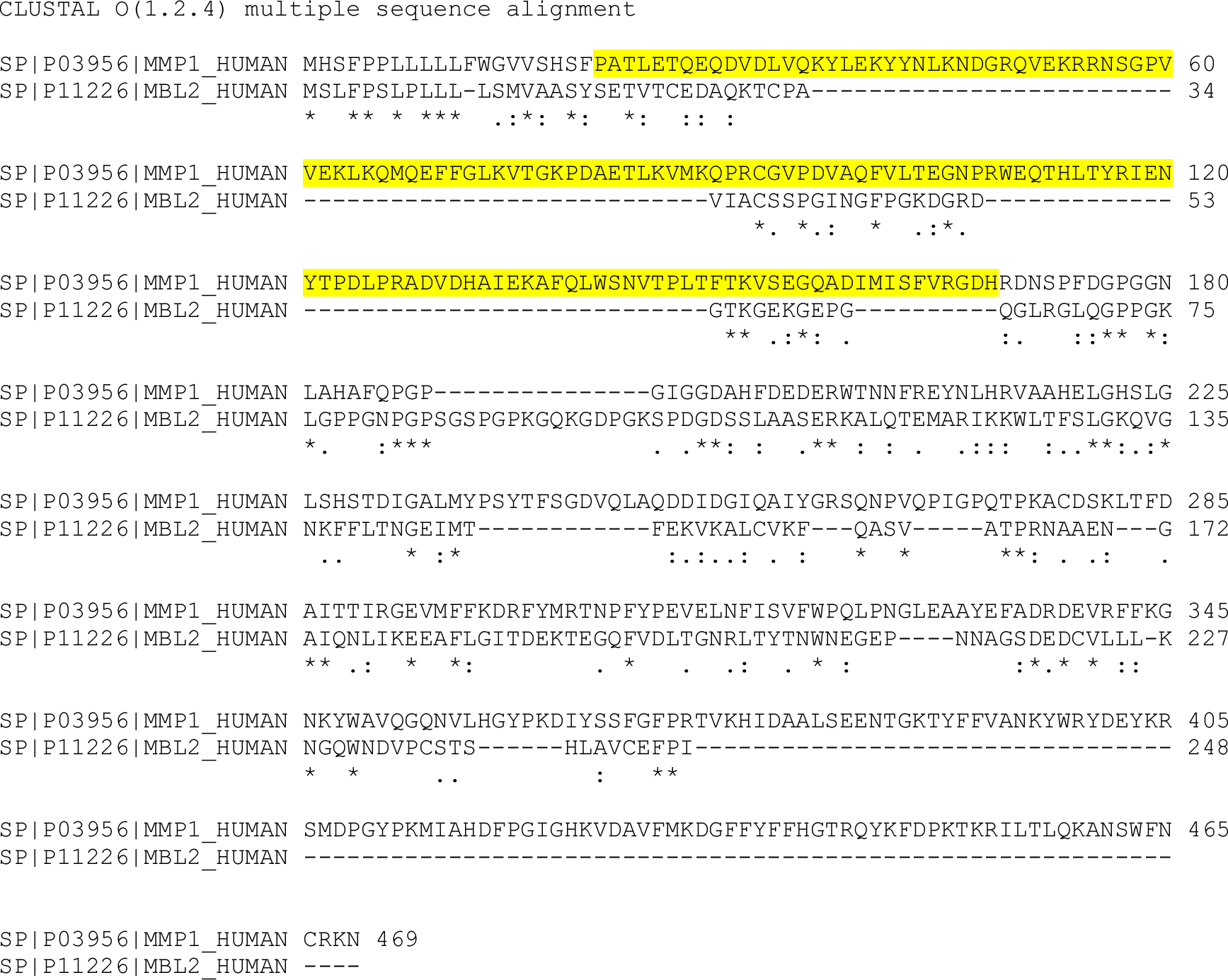
Sequence homology search of MMP1-MBL2 using CLUSTALO alignment revealed a 10.33 *%* homology. CLUSTAL. In Yellow: Amino acidic sequence used as antigen in the generation of the antibody(PrEST).

**Supplementary Figure 8.**
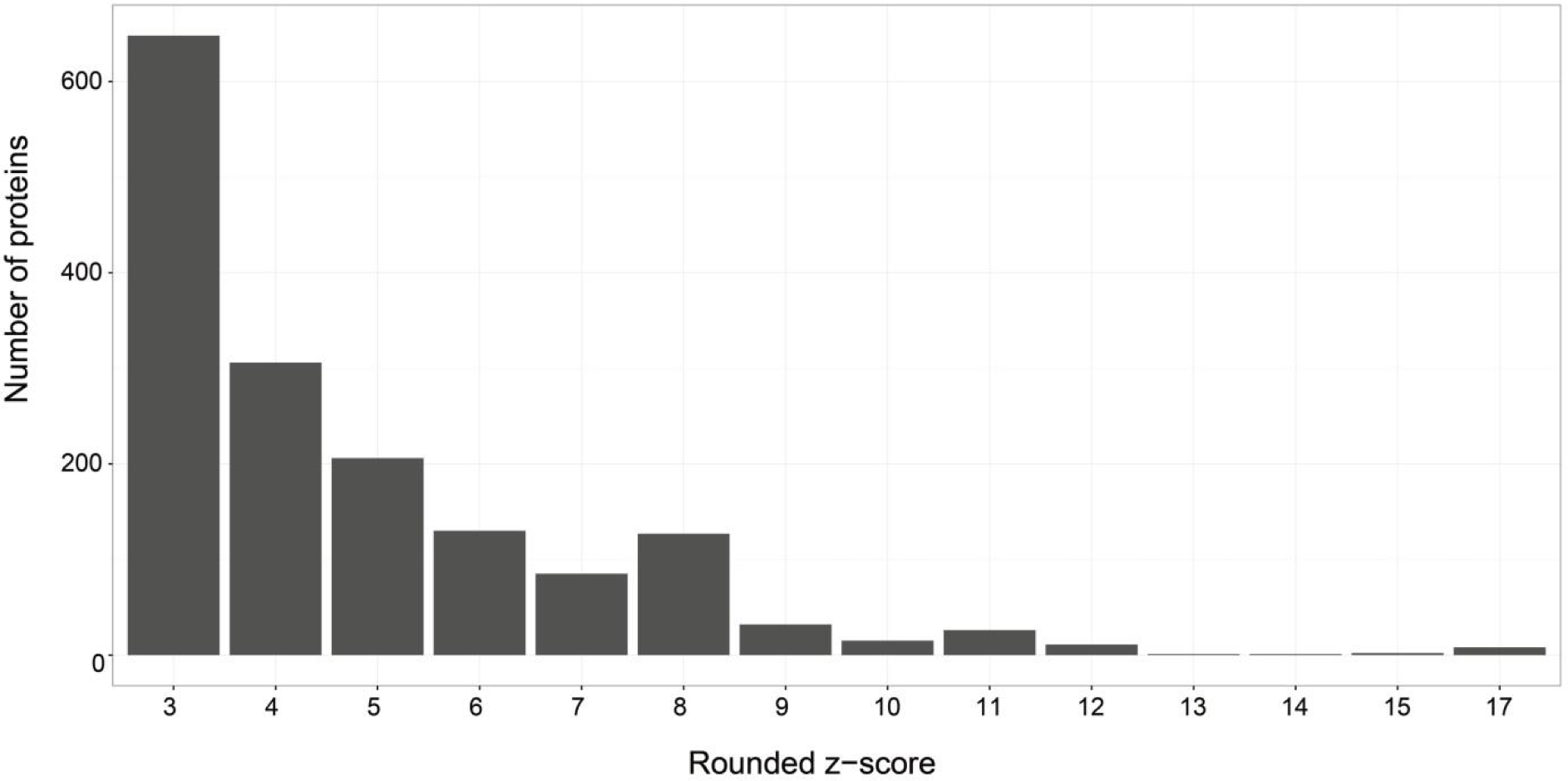
Bar plot representation of the number of proteins to which a z-score ≥ 3 was assigned. Values of z-scores were rounded.

**Supplementary Figure 9:**
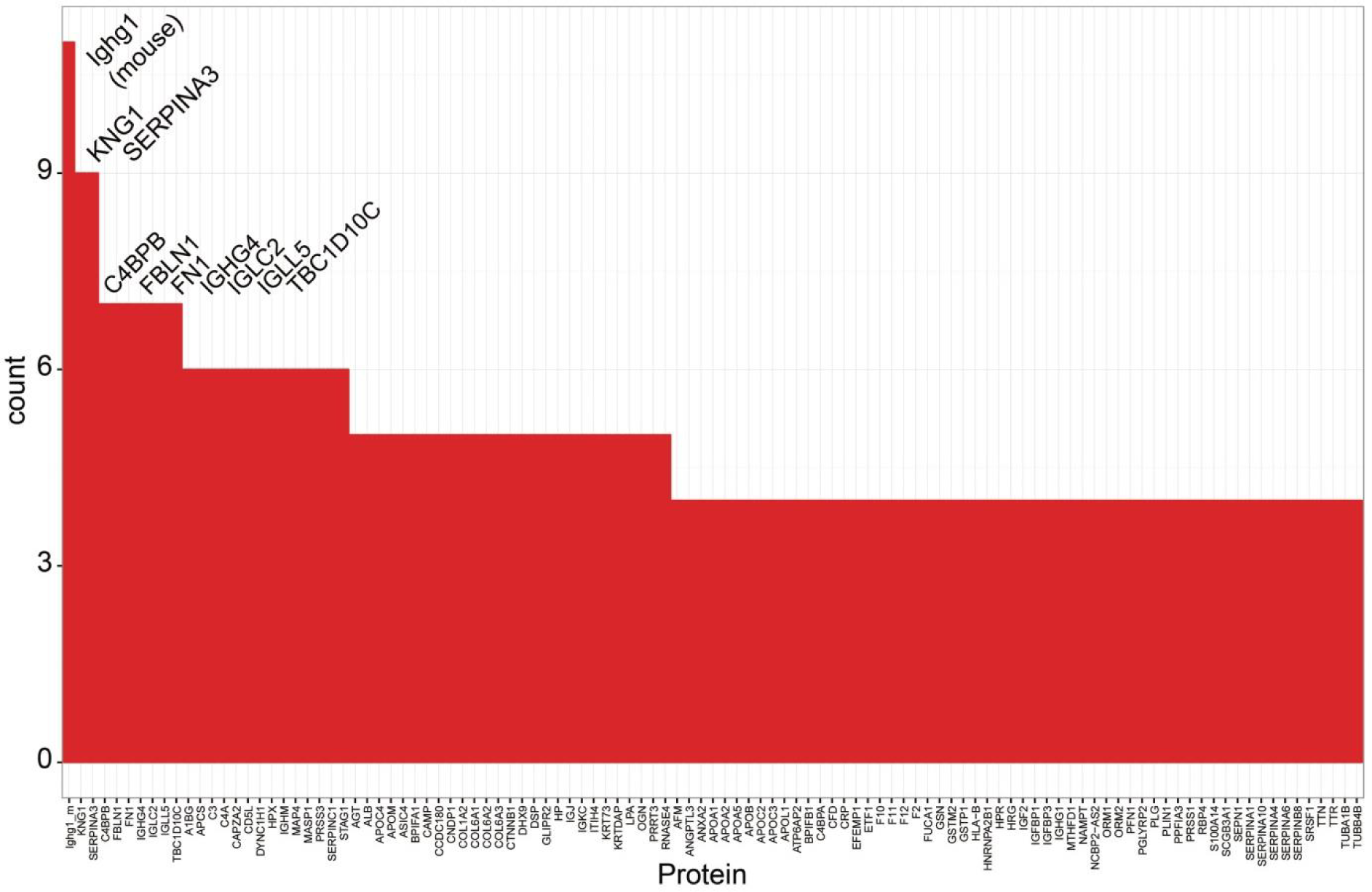
Bar plot representation of proteins identified co-targets for more than 3 antibodies.

**Supplementary Figure 10.**
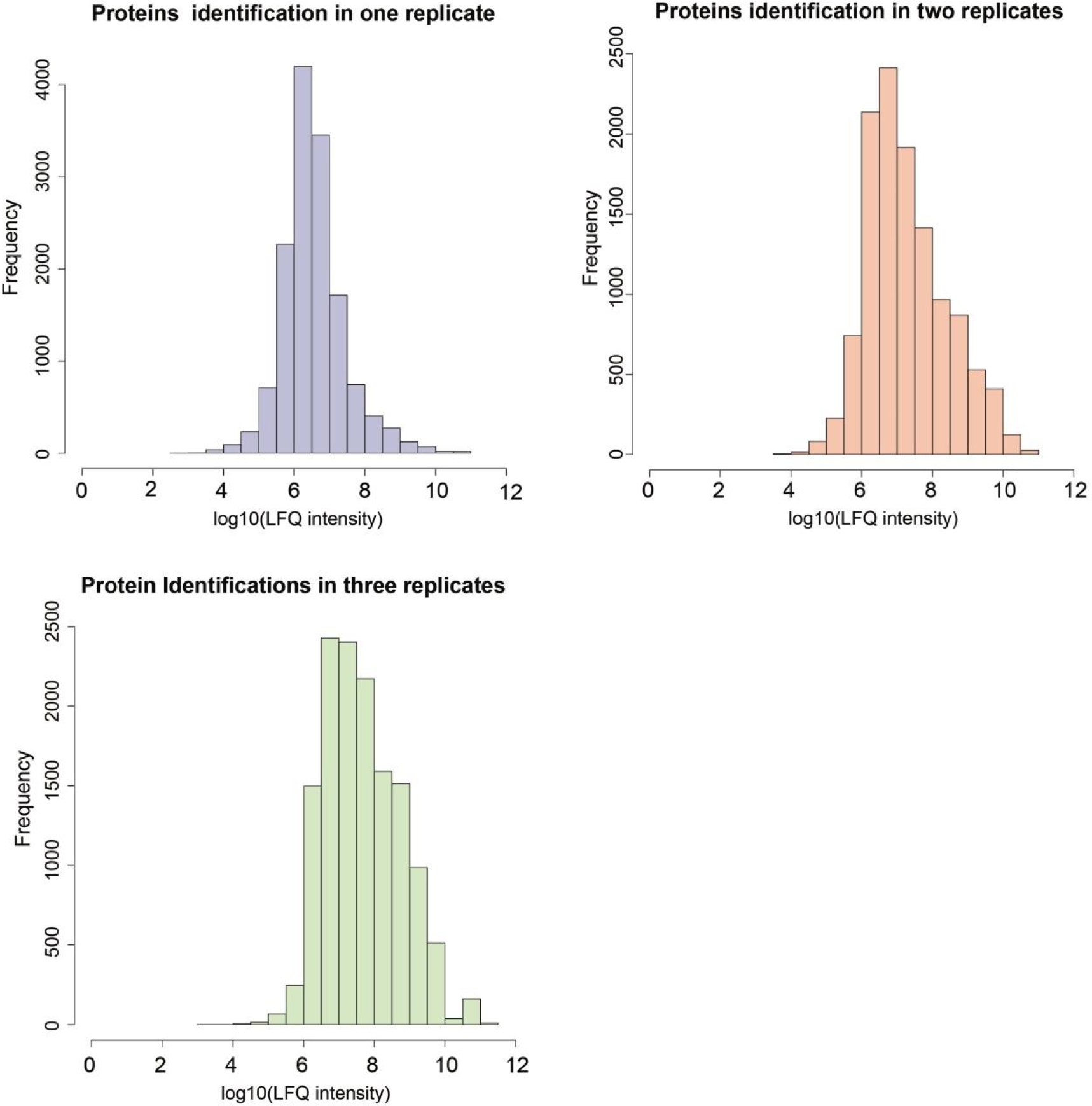
Defining a LFQ cut-off base on the Distribution of LFQ intensities associated to proteins identified (A) in a single; (B) duplicate and (C) in triplicate experiments.

**Supplementary Figure 11.**
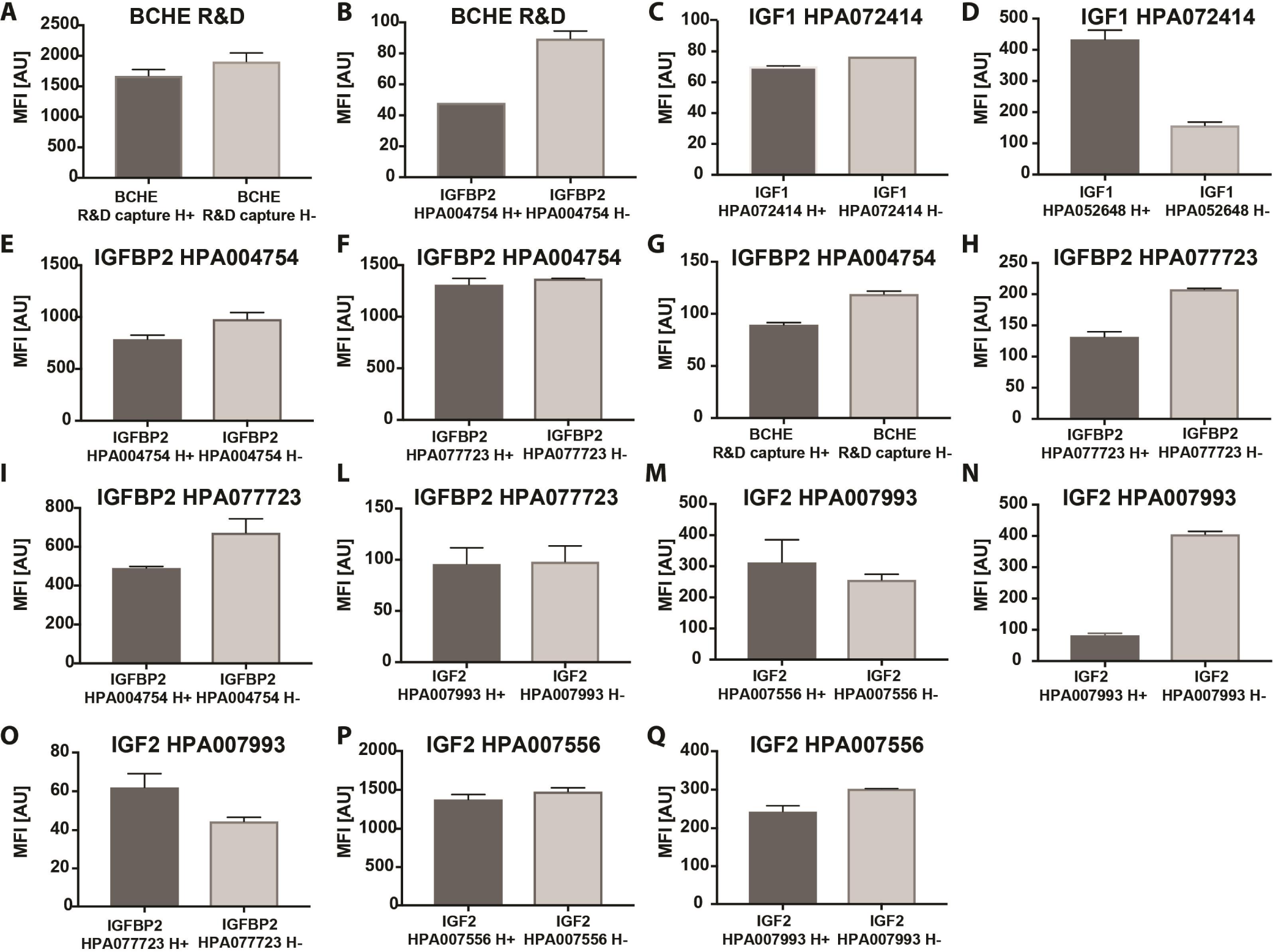
Combinations of antibodies were tested in plasma. The MFI values of the corresponding pair of antibodies indicated for the EC50 plasma concentration. Headers of the plots indicate the detection antibody and on the x axis is the capture antibody. The data calculated is based on the mean values (±SD) of duplicate dilution curves. Dark grey (H+, heated plasma), Light grey (H-, untreated plasma).

**Supplementary Table 1.**
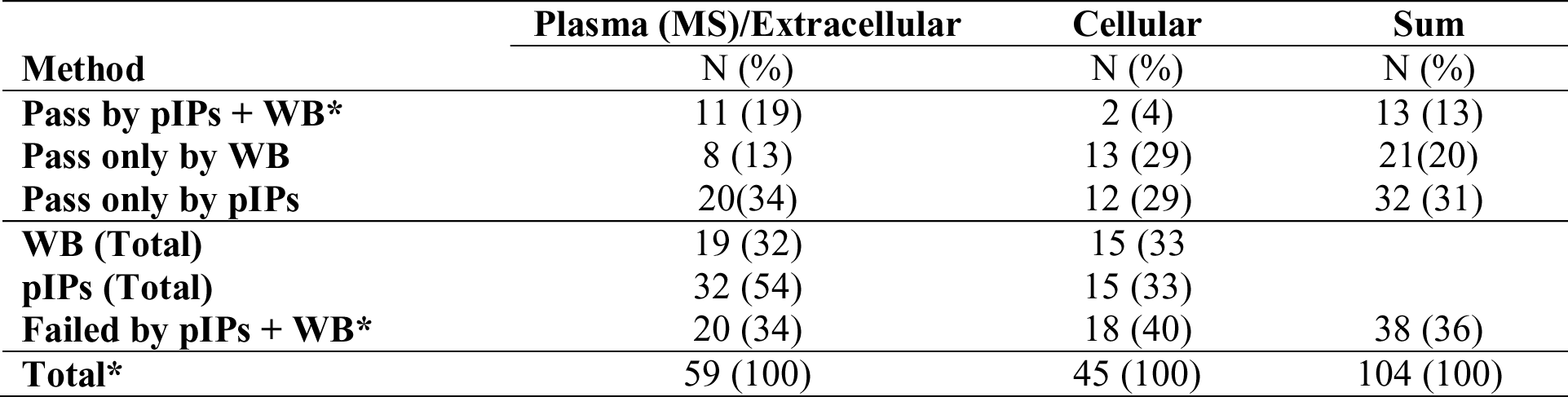
Comparison of validation rate by pIPs and WB for 104 antibodies assessed previously in plasma by WB. An antibody was considered passed by WB if scored 1,2 or 3, according to the Human Protein Atlas project.

## SUPPLEMENTARY EXCEL TABLE

**Description of content of each Sheet.**

- “Selected targets”: Description of proteins for which antibodies were selected. (See Supplementary References 23-108)
- “Protein Annotation (GO)”: Protein annotation using GO terms.
- “Antibodies experim. annotations”: Antibodies used in the study, annotations about experimental conditions and data of validation by pIP and WB
- “Frequencies of identification”: List of proteins identified in pIP experiments and their frequencies of identification in heat treated or not heat treated plasma.
- “Batches Kruskal Wallis Test”: p-values for differences in intensity levels detected the between batches
- “Go enrichment analysis”: Differential GO analysis using TOPP cluster between frequent (> 20%) and less frequently occuring proteins (< 20%).
- “z-score >2.5”: List proteins for which a z-score > 2.5 was calculated containing targets, off-targets and potential interactors.
- “List of Peptides”: Peptides identified in pIP assays for the expected target proteins and overlapping with the antigen used to generate the antibodies.
- “Experimental Batches”: description of major differences in the experimental conditions for the 6 experimental batches analyzed.
- “Reagents Catalogs_Lot_numbers”: List of catalog and lot numbers for affinity reagents.
- “Antibodies against same protein”: proteins enriched by antibodies raised against the same target protein, using same or different protein fragments.

## MATERIAL AND METHODS

### 2.1 Sample collection

Human EDTA plasma from a pool of individuals (50% females) was obtained from Seralab (Sera Laboratories International Ltd). Aliquots of plasma (0.5 mL) were stored in cryogenic vials at -80°C and thawed at 4°C before use.

### 2.2 Target selection

Information about target proteins their functions and involvement in diseases were collected through literature searches, Gene Ontology (GO, http://www.geneontology.org/), Human Protein Atlas (HPA, http://www.proteinatlas.org/), and Early Detection Research Network (EDRN, https://edrn.nci.nih.gov/). Plasma protein abundances were obtained from the 2017 built of the Plasma PeptideAtlas ([1], www.peptideatlas.org). Proteins were classified using the following terms: (i) Cellular or Extracellular, when the proteins appeared in one or more of the terms extracellular region (Go:0005576); extracellular space (Go:0005615); extracellular exosome (Go:0070062); proteinaceous extracellular matrix (Go:0005578); (see columns “GO CC Complete” and “Summary of GO CC”in the Excel sheet “Protein Annotation GO”). The list of antibodies and information was reported in Supplementary Excel Tables. The analysis included 157 antibodies. There were 15 monoclonal antibodies (10 from R&D Systems; 1 from HyTest Ltd.; 3 from Atlas Antibodies and 1 from SigmaAldrich) and144 polyclonal antibodies from the Human Protein Atlas [2]. In addition, normal rabbit IgG (Bethyl Laboratories), mouse IgG and rat IgG (both Santa Cruz Biotechnology) were included as controls. Catalog and lot numbers are listed in Supplementary Excel Tables. For antibodies obtained by the Human Protein Atlas, antibody IDs and lot numbers are the same.

### 2.3 Antibody coupling to magnetic beads

Covalent coupling of antibody to magnetic beads (MagPlex, Luminex Corp.) was performed as previously described [3]. Briefly beads were activated using sulfo-NHS (Sulfo-N-hydroxysulfosuccinimide, Thermo) and ethyl-carbodiimide (EDC, both Thermo). Then 1,6 μg antibodies diluted in MES buffer per 500 000 beads were incubated 2 h at room temperature, beads were washed and stored in blocking buffer at 4 °C.

### 2.4 Immunocapture-mass spectrometry

Aliquots of EDTA plasma (Seralab) were diluted 1:10 in assay buffer containing 0.5% w/v PVA (Sigma-Aldrich), 0.8% w/v PVP (Sigma-Aldrich), 0.1% w/v casein (Sigma-Aldrich) and 10 % of rabbit IgG (Bethyl Laboratories, Inc.). Samples undergoing heat treatment were incubated for 30 min at 56 °C in water bath, before being combined with beads and incubated overnight on a rotation shaker at 23°C. For the final assessment of 153 antibodies 100 μl of crude plasma and 1,6 μg of antibody couple to beads were applied in each incubation. On the next day, and using a magnetic bead handler (KingFisher™ Flex Magnetic Particle Processors, Thermo Scientific), beads were separated from the sample, washed with 0.03% Chaps in PBS and re-suspended in digestion buffer containing 50 mM ammonium bicarbonate (Sigma-Aldrich) and 0.25% sodium deoxycholate (Sigma-Aldrich). Proteins were reduced with 1 mM DTT (Sigma-Aldrich) at 56 °C for 30 min, and alkylated by iodoacetamide 4 mM (Sigma-Aldrich), at RT in the dark for 30 min. Alkylation was quenched adding 1 mM DTT. Proteins were digested using a mixture of Trypsin and LysC at 1:25 trypsin-to-protein ratio (Promega, USA) overnight at 37 °C. Enzyme inactivation and sodium deoxycholate precipitation was obtained adding 0.005% TFA. Peptides in the supernatant were then separated from beads, dried and re-suspended in solvent A containing 3% acetonitrile (ACN) and 0.1% formic acid (FA).

### LC-MS/MS

MS analysis was performed using a Q-Exactive HF (Thermo) operated in a data dependent mode, equipped with an Ultimate 3000 RSLC nanosystem, Dionex). Samples were injected into a C18 guard desalting column (Acclaim pepmap 100, 75 μm x 2 cm, nanoViper, P/N 164535, Thermo) and then into a 50 cm x 75μm ID Easy spray analytical column packed with 2μm C18 (EASY-Spray C18 P/N ES803, Thermo) for RPLC. Elution was performed in a linear gradient of Buffer B (90% ACN, 5% DMSO, 0.1% FA) from 3 % to 43% in 50 min at 250 nL/min. The proportion of Buffer B was increased stepwise to 45% in 5 min, then to 99% in 2 min, and then held for 10 minutes. DMSO was added Buffer A for the chromatography (90% water, 5% ACN, 5% DMSO, 0.1% FA). Full MS scan (300-1600 m/z) proceeded at resolution of 60,000. Precursors were isolated with a width of 2 m/z and listed for exclusion for 60 s. The top five most abundant ions were selected for higher energy collision dissociation (HCD). Single and unassigned charge states were rejected from precursor selection. In MS/MS, a max ion injection time of 250 ms and AGC target of 1E5 were applied.

### 2.7 Data analysis

Shotgun MS data search was performed on MaxQuant (v1.5.3.30) [4] using the integrated algorithm MaxLFQ. Spectra were search against a human protein database from Uniprot (accessed on 03/17/2016, Canonical and Isoforms, 20,198 hits customized adding sequences of immunoglobulins chain C from rabbit, rat and mouse, LysC (PSEAE) and Trypsin (PIG). Settings allowed for two missing cleavages, methionine oxidation and N-term acetylation as variable modification and cysteine carbamidomethylation as fixed modification. Fast LFQ and match between runs were applied, three minimum number of neighbors, and six average number of neighbors. All the 414 raw data files included in the analysis of 153 antibodies plus controls were analyzed in a single session, LFQ intensity values obtained were used for the following analysis. We considered as contaminants: proteins belonging to the list of contaminants in MaxQuant not belonging to Homo sapiens (Human), Ig gamma chain C region from Oryctolagus cuniculus (Rabbit)) because known to be in the dilution buffer andImmunoglobulin variable chains belonging to Homo sapiens (Human), and excluded from the z-score calculations. We considered missing values as missing not at random (MNAR) [5], but missing because of concentrations below the limit of detection (LOD). We therefore used min = 0 as minimum value detected of intensity (Single-value imputation approach). When calculating average and standard deviation for each protein identified over the population of all experiments, missing values of LFQ intensities were substituted to 1 to allow for log10-transformation before two ways hierarchical clustering and principal component analysis. For z-scores calculation, when duplicate and triplicate experiments were available, we considered only proteins identified in all replicates for further analyses, and calculated average of LFQ intensities Proteins were considered enriched when associated to a z-score ≥ 3. To visualize the enriched proteins for each antibody, z-scores and LFQ intensity values were use and proteins found above the set threshold were annotated accordingly. Raw data produced to assess experimental conditions were analyzed using MaxQuant but excluding the function for LFQ.

Data analysis and representation was performed on the environment for statistical computing and graphics R [6]. Alignments between protein and prEST sequences was performed using the Clustal Omega program available at EMBL-EBI [7]. GO enrichment system was performed using the PANTHER Classification System (http://pantherdb.org/). Comparison of GO terms was conducted using ToppCluster ([8]; https://toppcluster.cchmc.org/), regarding Bonferroni corrected p-values < 0.01 as significant.

### 2.8 Sandwich Immunoassay

The capture antibodies towards IGFBP2, IGF1, IGF2, DERA, BCHE, and rabbit-Immunoglobulin G (rIgG) and mouse-IgG, as negative controls (**Supplementary Excel Table**, sheet: “Reagents Lot numbers”), were covalently coupled to color-coded magnetic beads, A Suspension Bead Array (SBA) was generated and analyzed in Luminex Platform as previously described [9], using an in house protocol for labeling detection antibodies with biotin [10]. The antibodies were coupled to beads and labeled in order to test each different combination of capture and detection antibody pairs listed in the Supplementary Excel Table.

Plasma (EDTA Seralab, LOT#BRH1147432) was thawed on ice and centrifuged for 1 min at 2000 rpm, and diluted from 1:20 following 4-fold dilutions in PVX casein (PVXC) buffer 10% rIgG. The dilution series consisted of 6 points in duplicate and were heated at 56°C for 30 min. Then, the plasma was incubated with the SBA overnight. The same procedure was carried out with non-heated plasma dilution series.

The recombinant proteins used were IGFBP2 and IGF1 were a kind gift from Hanna Tegel and Johan Rockberg (AlbaNova University Center, KTH), IGF-II (R&D systems, catalog # 292-G2-050, lot DS2416011) and BCHE (DuoSet kit R&D systems, Catalog # DY6137-05, lot # 1387842). The dilution series of the proteins in buffer (PVXC 10% rIgG) consisted of 7 points prepared in duplicate and heated at 56°C for 30 min before incubation with the SBA overnight. Non-heated protein dilution series were also tested. The standard curves comprised different concentration ranges depending on the protein of interest. IGFBP2 and IGF2 were diluted in buffer from 500 ng/mL following 3-fold dilutions, IGF1 from 12000 pg/mL following 3-fold dilutions and BCHE 10000 pg/mL following 2-fold dilutions.

The detection antibodies were applied at 1μg/mL (HPA antibodies) or 25 ng/mL (BCHE R&D systems) for 90 min, and streptavidin-R-phycoerythrin (R-PE) conjugate (Life Technologies; SA10044) was used for the fluorescence read out in FlexMap3D (Luminex Corp.).

## SUPPLEMENTARY NOTES

**Supplementary note 1: Optimization of experimental conditions and quality control**

Factors such as target protein concentration and properties of ionization of the peptides of interest would ideally require optimized analysis for each antibody/target. Nevertheless, we began with developing a procedure applicable to a broad range of antibodies and target proteins. In order to set up optimal experimental conditions, we evaluated technical aspects such as volume of neat plasma andamount of antibody. As expected, increasing the volume of plasma, peptides belonging to less abundant proteins became detectable. For example interleukin 1 alpha (IL1A), with concentrations ~3 μg/mL in healthy human plasma [11], was detectable when 1 mL of plasma and 3.2 μg of antibody were applied (**Supplementary Figure 2A**). We established our protocol to enable the detection of proteins in a range of concentrations from μg/mL (C2) [12] to high pg/mL (KLK3) [13] [14-17] (**Supplementary Figure 2B**).

**Supplementary note 2: Batch effects**

We observed that independent IPs performed in the same batch clustered together (**Supplementary Figure 3A**). Parameters varying between independent batches IPs include different lots of reference plasma, trypsin, and analytical columns. Long-term drift in instrumental response, sample handling and sample heat-treatment may also add additional variability. We found that the main difference between the different assays was due to heat treatment of the samples. Indeed, despite experimental batches, assays using either heat treated or untreated plasma clustered together (Supplementary Figure 4A-B). For this reason, we decide to analyze IPs with heat treated and not heat-treated plasma separately, in order to compare the enrichment profiles from the IPs with similar background.

**Supplementary Figure 2A:**
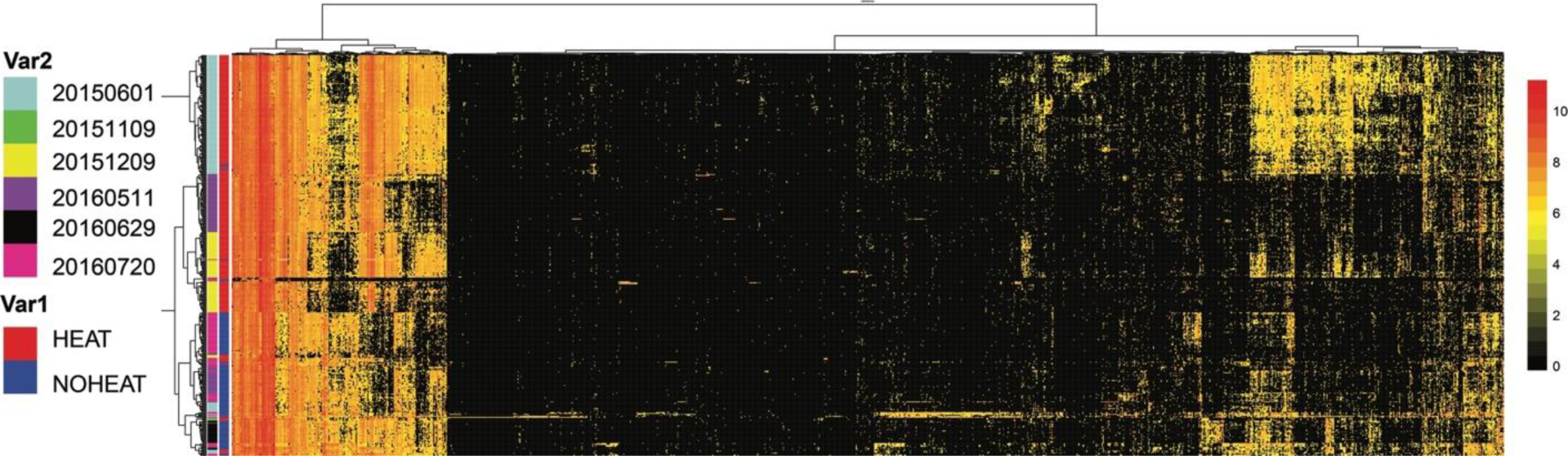
Two ways hierarchical clustering analysis of the proteins identified in 414 immunoprecipitations. Clustering distance: "euclidean”; Clustering method: Ward. Cluster lanes left to the heat map represent (1) different experimental batches and (2) sample heat treatment. (B) Representation of principal component analysis, dots colors highlight separated experimental batches.

**Supplementary Figure 2B:**
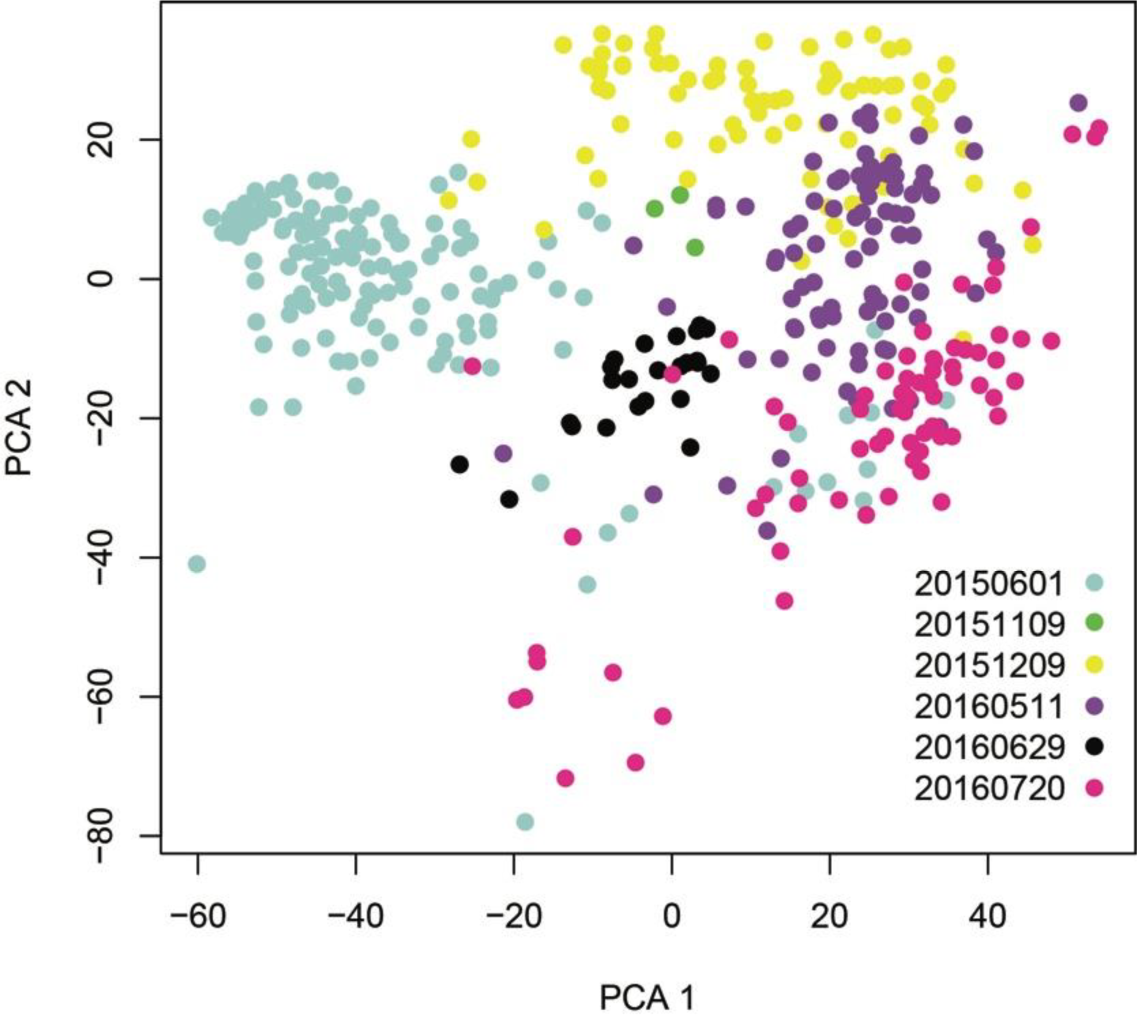
Representation of principal component analysis, dots colors highlight separated experimental batches.

**Supplementary note 3: Comparison of estimated plasma concentration and frequency**

We investigated the relation between the frequency of occurrence of 1313 proteins in untreated plasma (**Supplementary Figure 3A**) and heat-treated plasma (**Supplementary Figure 3B**) with the estimated plasma concentration from PeptideAtlas. There was a significant association (p-value <2.2e-16) for both sample types while 293/1313 = 22% of the proteins (blue dots) were not found with a concentration estimation in PeptideAtlas.

**Supplementary note 4: Investigation of passenger proteins**

We determined possible presence of antigens in the antibody solution. Antigens that were used during the affinity purification may have been co-eluted. Each antigen, denoted PrEST, consists of a tag (His_6_ and albumin binding protein ABP) and the region selected target protein [18]. To detect the presence of the antigens, we chose an antibody specific to tag region (1:60,000; [19]) and incubated antibody-coupled beads accordingly. Detection using an anti-IgY antibody carrying R-PE (1 μg/ml) we determined the cut-off as 10x SD + mean MFI obtained from beads carrying normal rabbit IgG. Out of 4,180 antibodies tested (See Supplementary Figure 4), 41 had been investigated for IP-MS. Among these, 11 revealed MFI above cut-off (135 AU). These included on binder of the ON-target category (HPA042270), nine CO-target binders (HPA002869, HPA005905, HPA019493, HPA030603, HPA045140, HPA045822, HPA062231, HPA067590, HPA070841) and one OFF-target binder (HPA057179).

**Supplementary note 5: Antibodies evaluation by protein arrays, WB and pIP.**

In plasma proteomics, current options for assessing the selectivity of antibodies can be offered by paired antibodies, protein arrays [20] or Western blot (WB) [21]. Even if the three methods are each valuable in evaluating antibodies quality, validation of a previous discovery or qualification prior to a planned assay should be performed by a method resembling as much as possible the experimental conditions of the previous or intended application.

For both protein arrays and WB, the setup is that a surplus of antibodies is diluted in a solution and applied onto supports that present the antigens. Hence, there is generally minimal competition for binding sites between potential on-and off-targets as compared to antibodies being immobilized and applied to a complex solution. For an example given by anti-IL6R binder, we found application dependent recognition for five different antibodies. All five were classified as target-specific using protein arrays, however only three detected IL6R in plasma (**Supplementary Excel Table**). It is consequently the composition of plasma, where 90% of protein content is assigned to 20 proteins, that poses a challenge for WB in terms of analytical resolution. In samples in which the concentrations of certain proteins dominate, the efficiency of how proteins migrate through the gel can be affected. This can influence the distribution of proteins in terms of their molecular mass and abundance; hence limits the amount of plasma loaded on the gel and so detectability of less abundant targets.

As first, we compared the classifications obtained here with existing scores of validation assigned by plasma WB to the same antibody’s Lot (HPA ID) within the Human Protein Atlas project [21]. Noteworthy, the comparison is based on different samples and the samples chosen for WB analysis were depleted of albumin and IgG prior use [22]. Nevertheless, we found that the assessment of 13 out of 104 antibodies (12%) provided supportive evidence by both methods. For antibodies raised against plasma proteins, the success rates for plasma IP (54%) was though higher than for plasma WB (32%) (**Supplementary Table 1**). When considering cellular proteins, the success rates were the same 33%. For WB however, uncertainty will remain unless other standards or comparative analyses are used to reference the detected bands. Until then, bands detected at the predicted molecular weight may still represent the recognition of an off-target molecule.

Consequently, pIP provides an unequivocal identification of the target and could elucidate ambiguous WB results (see C1orf64, CEP162, E2F7,and CCL16 **Supplementary Excel Table**, sheet: ”Antibodies_experim_annotation”). While WB may provide a more accessible technology for some labs, the information added by identification in MS will be required for an in-depth analysis of antibody selectivity. IP-MS can here be one option to enhance our understanding of binding to proteins, in particular for plasma.

**Supplementary Table 1.** Comparison of validation rate by pIPs and WB for 104 antibodies assessed previously in plasma by WB. An antibody was considered passed by WB if scored 1,2 or 3, according to the Human Protein Atlas project.

|  | **Plasma** | **Cellular** | **Sum** |
| --- | --- | --- | --- |
|  | **(MS)/Extracellular** |  |  |
| **Method** | **N (%)** | **N (%)** | **N (%)** |
| **Pass by pIPs + WB\*** | 11 (19) | 2 (4) | 13 (13) |
| **Pass only by WB** | 8 (13) | 13 (29) | 20(20) |
| **Pass only by pIPs** | 21(35) | 13 (29) | 34 (33) |
| **WB (Total)** | 19 (32) | 15 (33) |  |
| **pIPs (Total)** | 32 (54) | 15 (33) |  |
| **Failed by pIPs + WB\*** | 19 (33) | 17 (38) | 36 (35) |
| **Total\*** | 59 (100) | 45 (100) | 104 (100) |

**Supplementary note 7: Interaction and homology analysis**

